# Unifying single-cell annotations based on the Cell Ontology

**DOI:** 10.1101/810234

**Authors:** Sheng Wang, Angela Oliveira Pisco, Aaron McGeever, Maria Brbic, Marinka Zitnik, Spyros Darmanis, Jure Leskovec, Jim Karkanias, Russ B. Altman

## Abstract

Single cell technologies have rapidly generated an unprecedented amount of data that enables us to understand biological systems at single-cell resolution. However, joint analysis of datasets generated by independent labs remains challenging due to a lack of consistent terminology to describe cell types. Here, we present OnClass, an algorithm and accompanying software for automatically classifying cells into cell types part of the controlled vocabulary that forms the Cell Ontology. A key advantage of OnClass is its capability to classify cells into cell types not present in the training data because it uses the Cell Ontology graph to infer cell type relationships. Furthermore, OnClass can be used to identify marker genes for all the cell ontology categories, independently of whether the cells types are present or absent in the training data, suggesting that OnClass can be used not only as an annotation tool for single cell datasets but also as an algorithm to identify marker genes specific to each term of the Cell Ontology, offering the possibility of refining the Cell Ontology using a data-centric approach.

## Introduction

Single-cell RNA sequencing (scRNA-seq) has emerged as a powerful tool to generate comprehensive organismal atlases encompassing a wide range of organs and tissues^1–10^. One of the most important tasks in single-cell analysis is cell type annotation because all downstream analysis heavily rely on such information. This process that aims at characterizing and labeling groups of cells according to their gene expression is currently very inefficient due to the intense need for manual curation by a panel of tissue experts for each tissue and organ^11–17^. Recent efforts in scRNA-seq have produced an unprecedented large compendium of expert-curated cell type annotations, paving the way for scientists to better understand cellular diversity^3,18^. However, utilizing these cell type annotations is challenging due to the inconsistent terminology used to describe cell types collected by independent groups. This inconsistency will likely increase as more groups generate new datasets and more cell types and states are characterized, thus substantially preventing reproducible annotations and joint analysis of multiple datasets.

The Cell Ontology offers a controlled vocabulary for cell types and has been proposed as the basis for consistently annotating large-scale single-cell atlases^19–23^. A natural approach to addressing the inconsistent vocabulary challenge is then to build computational methods that automatically assign cells from different datasets to categories in the Cell Ontology. Ideally, these methods should be fully automated such that the results can be quickly updated as the Cell Ontology evolves.

However, assigning cells to terms (i.e., cell types) in the Cell Ontology has at least three challenges. First, although the Cell Ontology contains valuable hierarchical relationships among cell types, most of these cell type terms are not associated with marker genes which are crucial for cell type annotation. Second, even though supervised learning approaches can be used to predict Cell Ontology terms that have curated annotations, they are unable to classify cells to unseen terms (i.e., terms that do not have any annotated cells in the training data). This issue largely prevents us from fully understanding cellular diversity as more than 95% of cell types in the Cell Ontology are unseen even in the largest datasets. Throughout this paper, we refer to “**unseen Cell Ontology terms**” to describe cell types from the Cell Ontology that do not have any annotated cells in the training data. In contrast, we use “**seen Cell Ontology terms**” to denote cell types with some annotated cells in the training data. Third, as the Cell Ontology is not developed specifically for scRNA-seq, it likely misses new cell types and cell states and so certain cell type relationships might be inaccurate. Collectively, these challenges hinder progress towards comprehensive cell type annotation and cellular diversity understanding.

We developed Ontology-based Single Cell Classification (OnClass) to address these challenges. OnClass is able to automatically classify cells to any cell type as long as its corresponding term is captured in the Cell Ontology, even if this cell type does not have annotated cells in the training data. To achieve this, OnClass first infers similarities among all cell types according to their distances in the Cell Ontology graph. It then leverages these cell type similarities to classify cells into unseen Cell Ontology terms based on the annotated cells of other seen Cell Ontology terms. OnClass can thus classify cells into any Cell Ontology term and consider even the hardest case when a term has no cell annotations in the training data. OnClass is the first method that can classify cells into a specific cell type (rather than into a generic unassigned category as previous work did^11,12^) even when the training set does not have any annotated cells for such cell type. Furthermore, by projecting single cell transcriptomes and the Cell Ontology into the same low-dimensional space, OnClass advances other important applications, such as marker genes identification.

We evaluated OnClass on the Tabula Muris Senis dataset^18^, representing the existing largest effort of cell type characterization. We found that our method outperformed existing methods at annotating both seen and unseen Cell Ontology terms. We further demonstrated the ability of OnClass to transfer annotations to 26 other single-cell datasets and assign cells to the correct cell type even for cell types that were not part of the training data. Finally, we showed OnClass was able to accurately identify marker genes for seen Cell Ontology terms as well as unseen Cell Ontology terms. These OnClass referred marker genes achieved comparable performance to curated marker genes on cell type annotation, paving the way for creating an organism-wide molecular representation of cellular diversity.

## Results

### Overview of OnClass

The Cell Ontology is a controlled vocabulary that organized 2331 cell types anatomically derived into a hierarchy based on the “is_a” relation. In OnClass, we first constructed a graph of cell types based on the hierarchical “is_a” relationship in the Cell Ontology and embedded these cell types into a low-dimensional space where similar cell types were close to each other^24,25^ (**Supplementary Note 1, Supplementary Fig. 1**). Single cell transcriptomes were then projected into this low-dimensional space by finding a nonlinear transformation that projected each annotated cell to the region of its cell type. Unannotated cells were also projected into this low-dimensional space using the same nonlinear transformation and annotated to the cell type corresponds to the region in which it lies. Importantly, such a procedure enables us to classify cells to unseen Cell Ontology terms based on their regions in the low-dimensional space. In addition to cell type annotation, OnClass used this low-dimensional space for other applications, including marker genes identification **(Fig. 1a)**. OnClass is Python-based open source package available at https://github.com/wangshenguiuc/OnClass. Our implementation can take as input a wide range of formats of the input gene expression matrix. It is able to consider any cell type similarity between the hierarchical structure of the Cell ontology used in this paper. Moreover, we provide a pre-trained model that is trained on TMS and can predict cell types for millions of cells in a few minutes on a modern laptop.

**Fig. 1.**
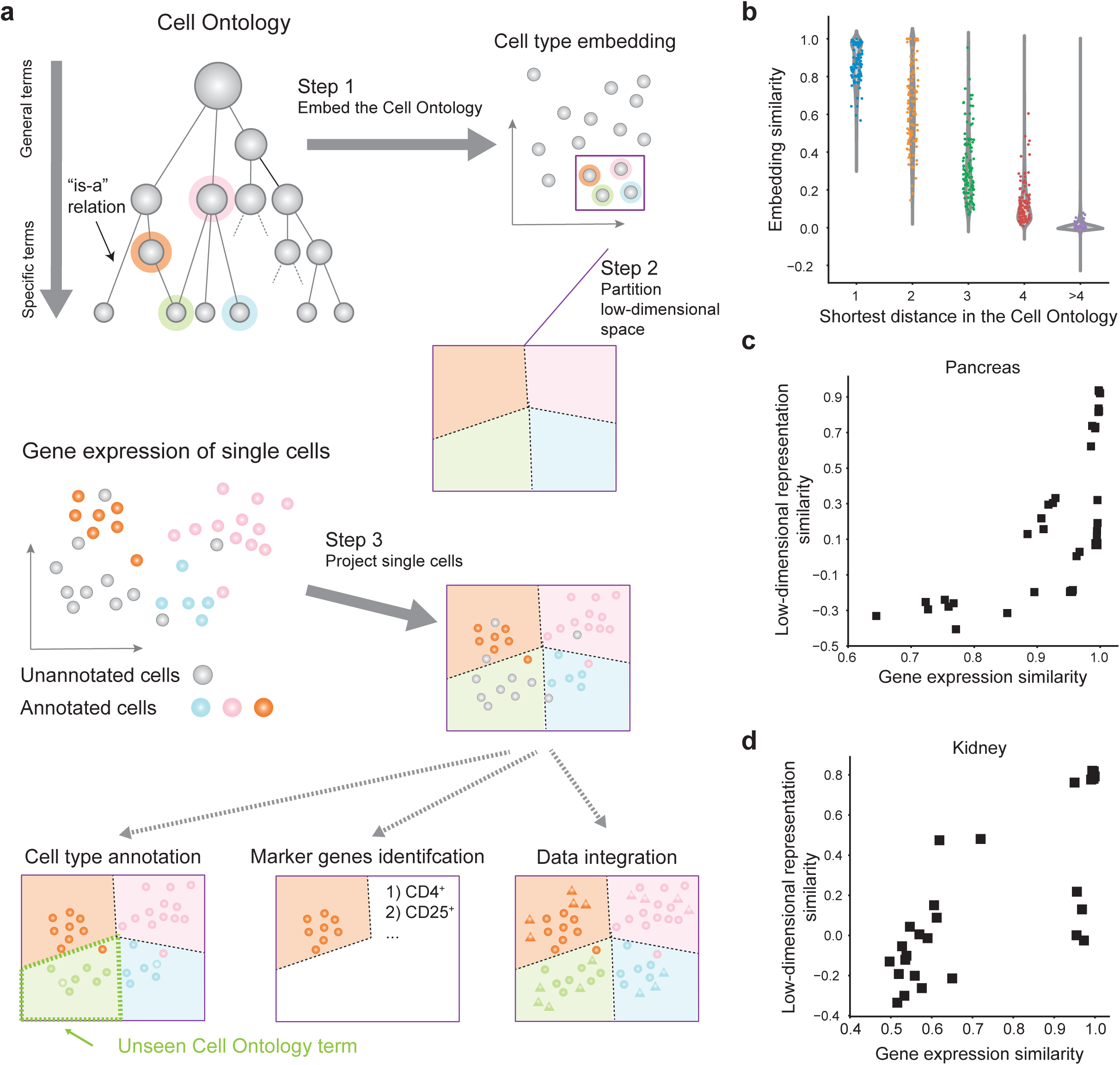
**a**, Flow chart of OnClass. The Cell Ontology is used to embed cell types into a low-dimensional space. OnClass then partitions this low-dimensional space into multiple regions, each corresponding to a cell type. Cells are then projected into this space by reducing the dimensionality of the gene expression matrix. These boundaries can then be used to annotate cell type, identify marker genes and integrate datasets. **b**, Violin plot showing the correspondence between the location of each cell type’s nearest neighbor in the Cell Ontology and the embedding similarity. The nearest neighbor of each cell type is calculated by using the cosine distance between cell type embeddings. **c,d** Scatter plots showing the correlations between the embedding-based cell type similarity and the gene expression-based cell type similarity in pancreas (c) and kidney (d).

### Cell type embeddings reflect cell type similarity

Since OnClass annotated cells even to previously unseen Cell Ontology terms according to the annotated cells from other Cell Ontology terms, its performance greatly relied on the quality of cell type embeddings. High-quality cell type embeddings should place cell types with similar gene expression profiles closely in the low-dimensional space, and can thus be used as good features for classification. Therefore, we first verified the merit of our approach by comparing three types of cell type similarities: the Cell Ontology structure-based similarity, the embedding-based similarity, and the gene expression-based similarity (**Methods**). We first observed that the embedding-based similarity was strongly correlated with the Cell Ontology structure-based similarity **(Fig. 1b)**. For example, the average embedding-based similarity of direct neighbors in the Cell Ontology graph was 0.86, which was 42% and 183% higher than the average embedding-based similarity of two-hop neighbors and three-hop neighbors. For cell types that are more than four-hop away in the Cell Ontology, the average embedding-based similarity was less than 0.01. Next, we examined whether cell types with similar embeddings would have similar gene expression profiles by comparing the embedding-based similarity and the gene expression-based similarity. Using a collection of annotated cells as the benchmark, we observed strong correlations between these two types of similarities. For instance, the correlation between the gene expression-based similarity and the embedding-based similarity was 0.70 (p-value < 1e-10) in pancreas and 0.77 (p-value < 1e-11) in kidney (**Fig. 1c,d**). The strong correlation between these two types of similarities demonstrated the high-quality of cell type embeddings and further suggested the possibility to annotate unseen Cell Ontology terms by using the knowledge from other similar and seen Cell Ontology terms. Unfortunately, none of the existing cell type annotation methods integrates with the Cell Ontology. OnClass’s ability to annotate cells with any cell type in the Cell Ontology led us to consider whether we could improve cell type annotation on large and diverse collections of scRNA-seq datasets.

### Improved cell type annotation using Cell Ontology

We ran OnClass on the Tabula Muris Senis (TMS) dataset^18^. To investigate the effect of unseen Cell Ontology terms, we split cells into test and training across different proportions of seen Cell Ontology terms in the test set. Overall, we observed that OnClass led to a substantial improvement in comparison to existing approaches (**Fig. 2a-d**). We first examined the ability of OnClass to identify cells belonging to a given Cell Ontology term. We observed that OnClass significantly outperformed all existing approaches in terms of AUROC on all proportions of seen Cell Ontology terms (**Fig. 2a**). Even when only half of Cell Ontology terms were observed in the training data, OnClass still achieved an AUROC of 0.87, while AUROCs of existing methods were all below 0.72. Next, we investigated whether OnClass could accurately predict the Cell Ontology term for a given cell. In a simpler setting where we combined all unseen Cell Ontology terms as a generic “unseen” class, OnClass outperformed existing methods in terms of Cohen’s Kappa statistic (i.e., balanced accuracy) from 10% to 90% of seen Cell Ontology terms (**Fig. 2b**). We found that the improvement of OnClass was more prominent with the increasing proportion of unseen Cell Ontology terms. We next evaluated a more challenging setting where unseen Cell Ontology terms were no longer combined and a prediction was deemed as correct only if the cell was assigned to the specific correct term, even if it is an unseen Cell Ontology term. By using Accuracy@3 and Accuracy@5 to quantify the performance, we observed significant improvement of OnClass in comparison to existing methods (**Fig. 2c,d**). For example, when 30% of Cell Ontology terms were unseen in the training data, OnClass obtained 0.45 Accuracy@3 and 0.55 Accuracy@5, while none of the existing approaches obtained accuracy greater than 0.3. Again, the improvement of OnClass was larger with more unseen Cell Ontology terms, indicating the advantage of using the Cell Ontology to transfer annotations from seen Cell Ontology terms to unseen Cell Ontology terms. To further demonstrate the importance of using the Cell Ontology, we found that the performance of OnClass substantially decreased by adding random noise to nodes (**Supplementary Fig. 2a**) and edges (**Supplementary Fig. 2b**) in the Cell Ontology. Notably, even though TMS had one of the most diverse and largest numbers of cell types, it still only covered less than 5% of all cell types in the Cell Ontology. Therefore, we anticipate that OnClass will be even more useful as more single cell RNA-seq datasets become available that contain transcriptomes for unobserved cell types in TMS.

**Fig. 2.**
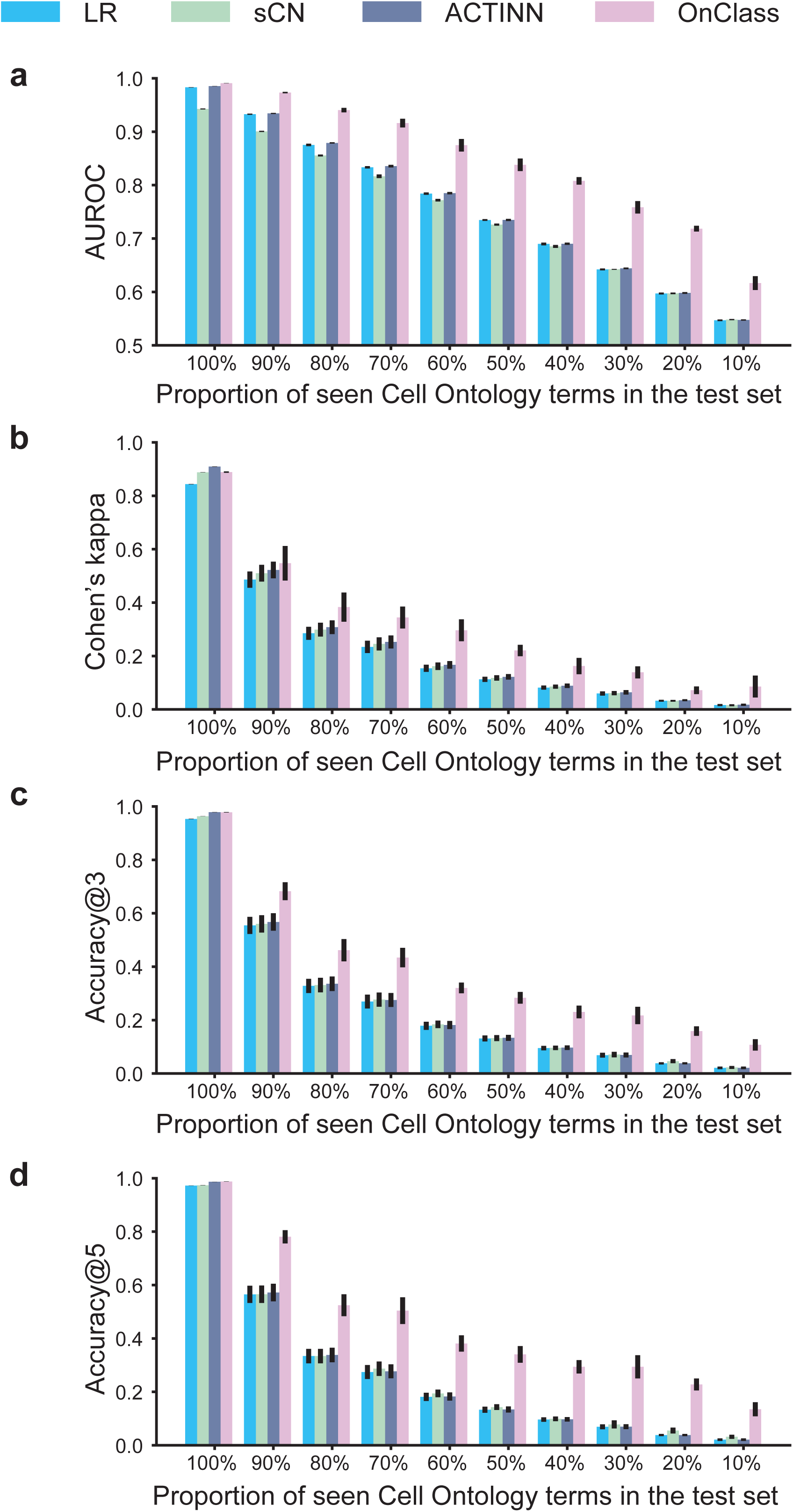
**a-d** Bar plots comparing OnClass and existing methods in terms of AUROC (a), Cohen’s Kappa (b), Accuracy@3 (c) and Accuracy@5 (d). x-axis shows the proportion of seen Cell Ontology terms in the test data.

### Annotating unseen Cell Ontology terms

We then examined the performance of OnClass in the more challenging case of annotating unseen Cell Ontology terms, which cannot be accomplished by any existing methods. Although recent efforts were able to classify cells into a generic “unknown” type^11,12^, they could neither break down this new type into detailed cell types nor attach it to a specific cell type term. To enable better comparison between OnClass and these approaches, we decided to extend existing approaches by classifying “unknown” type cells to the nearest cell type in the Cell Ontology (**Methods**). We studied the performance of OnClass by using an increasing number of seen Cell Ontology terms as the training data and all cells in the test data belonged to the remaining unseen Cell Ontology terms. We observed significant improvement with OnClass across different proportions of seen Cell Ontology terms. For instance, when using 60% of seen Cell Ontology terms as the training data (**Fig. 3a**), OnClass obtained an AUROC of 0.73. Even when only using 20% of cell types as the training data, OnClass still obtained an AUROC of 0.68. On a randomly selected set of 9 new unseen terms, OnClass was able to accurately classify 81% of cells (**Fig. 3b-d**). On a larger set of 21 unseen terms, OnClass still accurately classified 58% of cells (**Fig. 3h-j**). We showed the comparison of OnClass annotation and ground truth annotation in **Fig. 3b-j**. We found that OnClass was able to accurately classify a majority of cell types, including rare cell types. For those cells that were not accurately annotated, we found that the term assigned by OnClass was indeed biologically related to the ground truth Cell Ontology term. When evaluating OnClass for each tissue separately, we also observed good AUROCs from 0.84 to 0.93 on 21 tissues, with an average AUROC of 0.87 (**Supplementary Figs. 3-22**). As expert annotation can be imperfect and mostly limited to familiar cell types, OnClass can correct these false positives and broaden expert knowledge.

**Fig. 3.**
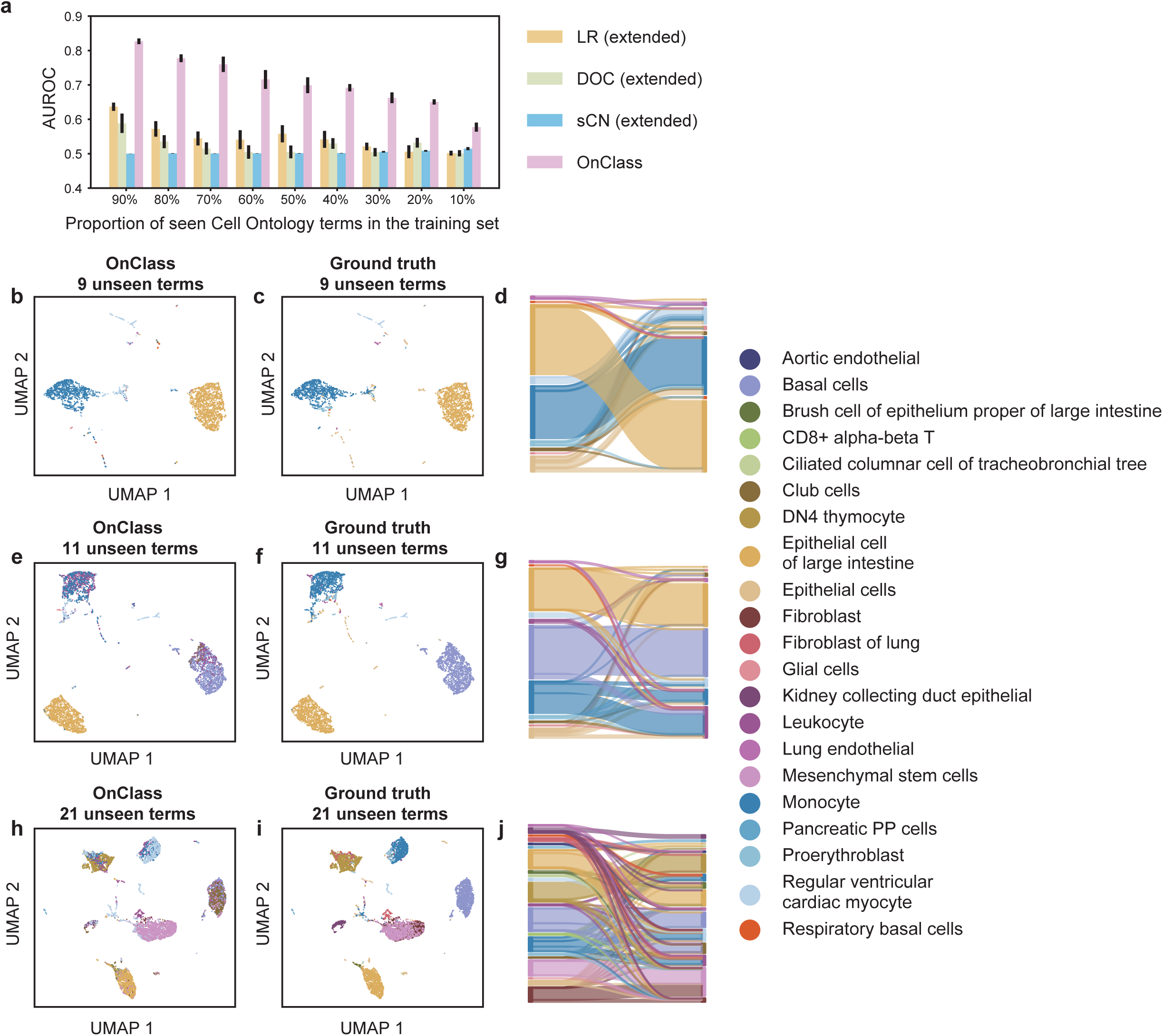
**a**, Bar plot comparing OnClass and existing methods for different proportions of seen Cell Ontology terms. x-axis shows the proportion of seen Cell Ontology terms in the training data and y-axis shows the AUROC. **b,c,e,f,h,i**, 2-D UMAP showing the predicted Cell Ontology terms of OnClass (b, e, h) and ground truth labels (c, f, i) for 9 unseen Cell Ontology terms (b, c), 11 unseen Cell Ontology terms (e, f), and 21 unseen Cell Ontology terms (h, i). The same color between OnClass predicted labels and ground truth labels means correct annotation. **d,g,j**, Sankey diagrams of the resulting mapping between predicted labels (left) to ground truth labels (right) for 9 unseen Cell Ontology terms (d), 11 unseen Cell Ontology terms (g), and 21 unseen Cell Ontology terms (j).

We next examined the robustness and applicability of OnClass by using it to annotate diverse datasets across animals, technologies, and organs. In particular, we used all cells in TMS to train OnClass and then classified 105,476 cells collected from 26 single-cell datasets (26-datasets) representing 9 technologies and 11 studies (**Methods**). We observed an average AUROC of 0.75 for these 26 datasets. Among all 10 cell types, OnClass obtained an AUROC greater than 0.8 for 5 of them (**Fig. 4a**). For B cell and macrophage that have annotated cells in TMS, OnClass obtained AUROCs of 0.99 and 0.97, respectively (**Fig. 4b, c**). More importantly, for cell types with no annotated cells in TMS, OnClass still achieved relatively high AUROCs (0.85 for CD14^+^ monocytes cell, 0.85 for CD56^+^ natural killer cell, and 0.81 for regulatory T cell), indicating its ability to accurately annotate and discover new cell types (**Fig. 4d-f**). Furthermore, the predicted cell type annotations can be used as features to cluster cells from different datasets. We used the predicted cell type annotations to combine these 26 datasets following the same procedure as previous work^26^. We observed good performance by using OnClass, where cells were clustered based on cell types rather than artifacts related to platforms (**Fig. 4g**). We further quantified the performance using the silhouette coefficient^27^ and observed a significant improvement in comparison to the state-of-the-art data integration approach Scanorama^26^ (**Fig. 4h**), indicating OnClass’s robustness to annotating cells from different batches and datasets.

**Fig. 4.**
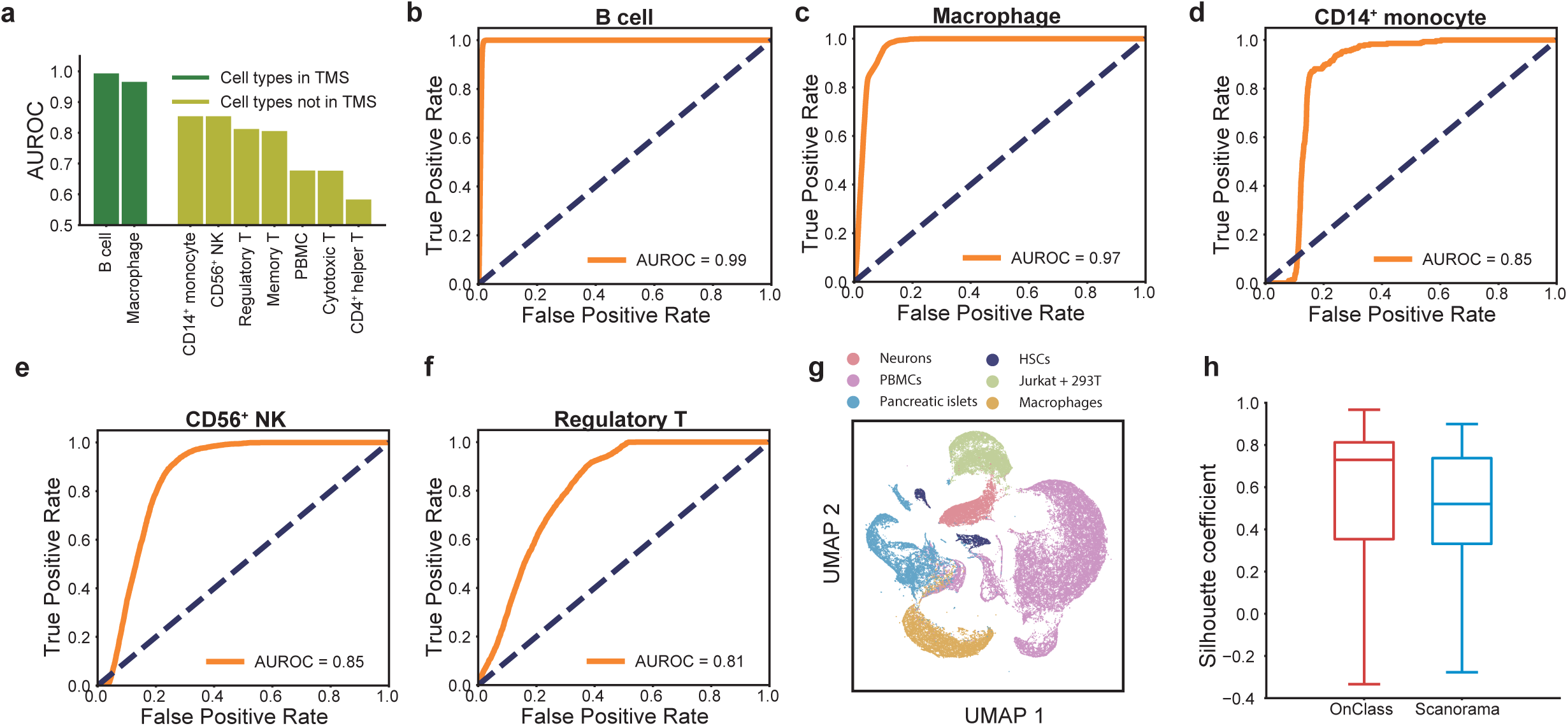
**a**, Bar plot showing the AUROC of OnClass on 9 cell types, including 2 present in TMS (green) and 7 not (yellow). **b-f** AUROC plots of OnClass’s prediction for five cell types: B cell (b), macrophage (c), CD14^+^ monocyte cell (d), CD56^+^ NK cell (e) and regulatory T cell (f). **g**, 2-D UMAP showing the 26 datasets and 6 cell types. **h**, Box plot showing the comparison between OnClass and Scanorama in terms of silhouette coefficient.

### Identifying marker genes for unseen Cell Ontology terms

Given the accurate annotation of both seen and unseen Cell Ontology terms, we were then interested in using OnClass to identify marker genes for the Cell Ontology terms. Marker genes are the key to expert curation but the existing knowledge is incomplete and limited extensively studied cell types. Here, we used OnClass to identify marker genes for both seen and unseen Cell Ontology terms in TMS (**Fig. 5a**). OnClass was able to identify the correct marker genes for 64% of seen Cell Ontology terms within the top 10 candidate genes in the predicted marker gene list. More importantly, since OnClass did not require any annotated cells to identify marker genes, it was able to find marker genes for unseen Cell Ontology terms as well. For example, OnClass identified the correct marker genes for 39% of unseen Cell Ontology terms within the top 10 candidate genes. We incorporated these OnClass-referred maker genes (**Supplementary Table 1**) and functions enriched with these marker genes (**Supplementary Table 2**) into our provisional Cell Ontology, in the hope of facilitating future expert curation. This data is easily accessible through our portal http://onclass.ds.czbiohub.org. Although these marker genes are by no means a completely accurate representation of cell type features, they are the first attempt at creating a comprehensive knowledge base of marker genes representative of the entire cellular diversity.

**Fig. 5.**
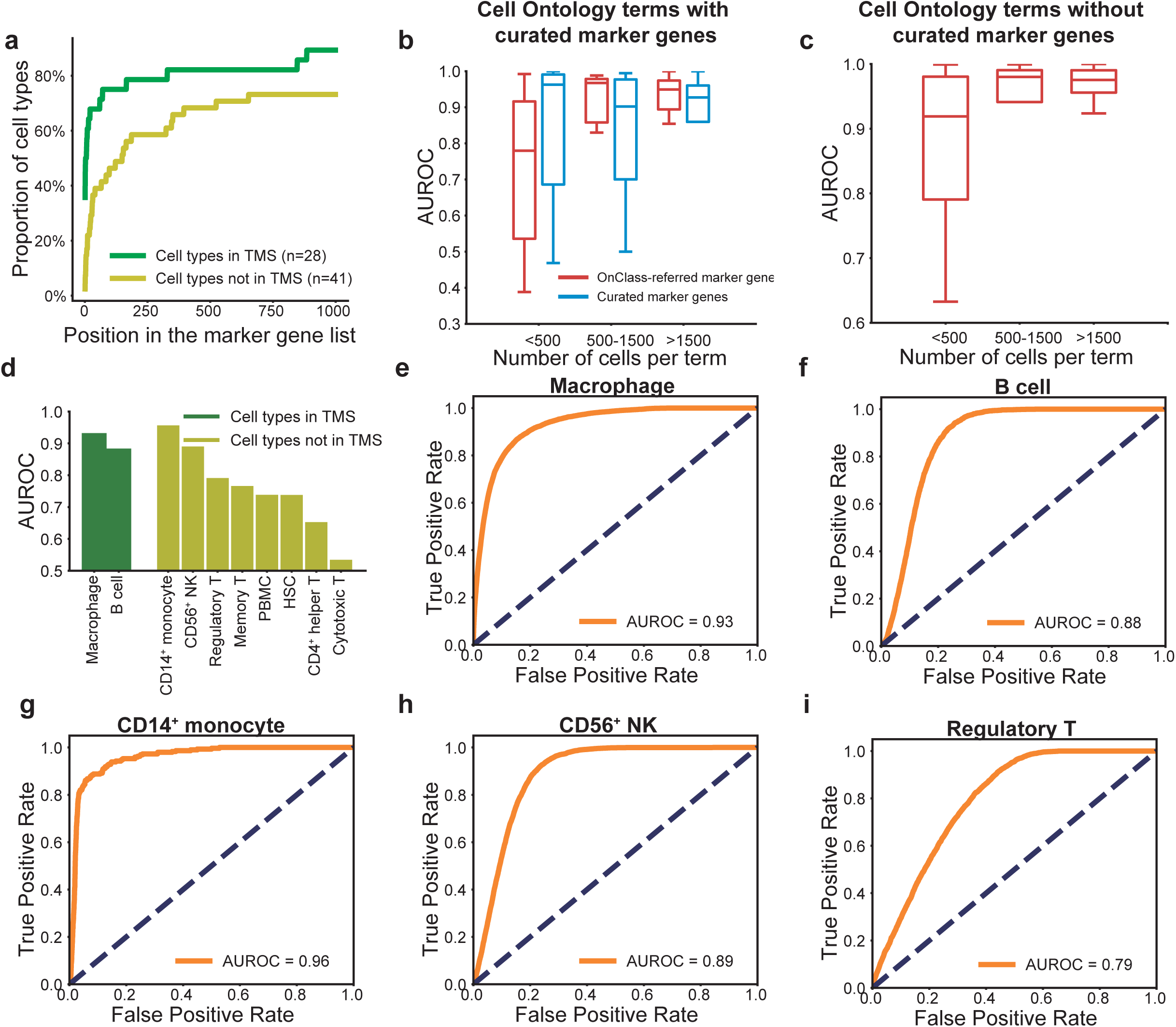
**a**, Plot showing the proportion of cell types out of the ones present (green) or not (yellow) in TMS for which OnClass can identify the marker genes in the top *k* genes out of 23,437 genes. *k* is shown in the x-axis and corresponds to the position in the marker gene list sorted by p-value. **b**, Boxplot showing the cell type annotation performance of using OnClass-referred marker genes (red) and curated marker genes (blue) in terms of AUROC. x-axis shows the number of cells per Cell Ontology term. **c**, Boxplot showing the cell type annotation performance of using OnClass-referred marker genes in terms of AUROC. Only Cell Ontology terms that have no curated marker genes are shown here. x-axis shows the number of cells per Cell Ontology term. **d**, Bar plot showing the AUROC of OnClass for 10 cell types, including 2 present in TMS (green) and 8 not (yellow). **e-i** AUROC plots of OnClass’s prediction for five cell types: macrophage (e), B cell (f), CD14^+^ monocyte cell (g), CD56^+^ NK cell (h), and regulatory T cell (i).

Finally, we sought to examine whether OnClass-referred marker genes could be used to accurately annotate cells. We first used all FACS cells in TMS to identify marker genes and then used these marker genes to annotate droplet cells in TMS. We found that the performance of using OnClass-referred marker genes was substantially better than using curated marker genes for Cell Ontology terms with more than 500 individual cells that were annotated in such category. For example, OnClass-referred marker genes achieved 0.98 AUROC, whereas curated marker genes achieved 0.90 AUROC for Cell Ontology terms with more than 500 and less than 1500 cells (**Fig. 5b**). For rare cell types, the performance of OnClass-referred marker genes was comparable to curated marker genes (**Fig. 5b**). Furthermore, for those Cell Ontology terms that have no curated marker genes, OnClass-referred marker genes also achieved accurate cell type annotation performance (**Fig. 5c**). We found that the performance of OnClass depends on the number of annotated cells and so as more data becomes available, we anticipate substantial improvement at the level of identifying robust and accurate marker genes. To assess the robustness of these marker genes, we next used these TMS-derived marker genes to classify 26-datasets. Among all the 10 cell types, 8 of them achieved AUROCs larger than 0.7 and 4 of them achieved AUROCs larger than 0.8 (**Fig. 5d-i**). Even for unseen Cell Ontology terms, OnClass still obtained a desirable performance (**Fig. 5g-i**). Notably, when comparing the performance with a supervised classifier, we found that using marker genes could achieve better results on several cell types (e.g., CD14^+^ monocyte cells) (**Fig. 4a, Fig. 5a**). Although supervised models are more expressive, they are also prone to overfitting. In contrast, marker genes are not only interpretable but also more robust to noise, thus enabling accurate annotation of new cells.

## Discussion

Ever since the emergence of scRNA-Seq, cell type annotation is a key step in single-cell data analyses. As more cell types are discovered and expected to be discovered, recent efforts have focused on classifying cells into existing labels or a generic unseen cell type^11,12^. Despite encouraging results based on these approaches, these methods fail to provide meaningful information specific to the cell types that are not part of the training sets. In contrast, our method takes an important step forward by mapping each cell to the Cell Ontology, leading to accurate annotations of cells with unseen Cell Ontology terms, which cannot be achieved by any existing methods. Conceptually and methodologically, this is substantially different from existing methods in the sense that our method explicitly leverages hierarchical cell-to-cell relationships to directly classify cells into any cell type within the Cell Ontology.

While our method leverages the Cell Ontology to classify unseen cell types, it is inspired by recent progress in single cell dataset integration approaches^26,28^. In the state-of-the-art single cell integration frameworks, datasets from different technologies are aligned in the same low-dimensional space by using mutual nearest neighbors as anchors to connect them. Indeed, our method can be considered to be aligning the Cell Ontology to the gene expression matrix by using known annotations as anchors. The key novelty of our method comes from effectively embedding cell types based on the hierarchical structure of the Cell Ontology and dividing the low-dimensional space into regions to enable assignment of cells to unseen Cell Ontology terms. We therefore expect our method to be of broad use to the community of cell biologists and computational biologists who are dwelling with the hard problem of identifying the cell populations present in each dataset.

---

**Supplementary Fig. 1**. Flowchart of the cell type embedding process. The Cell Ontology graph is constructed based on the “is_a” relation in the Cell Ontology. Random walk with restart is performed on the graph, restarting from each node. An equilibrium distribution is calculated for restarting from each node. These distributions are then concatenated and then reduced into a low-dimensional space.

**Supplementary Fig. 2a,b**, Plot of adding random noise on nodes (a) and edges (b) into the Cell Ontology graph. X-axis the ratio of random noise and y-axis is the AUROC.

**Supplementary Figs. 3-23**, Plot of the Cell Ontology of cell types that have annotated cells in aorta, BAT, brain myeloid, brain non-myeloid, diaphragm, GAT, heart, kidney, large intestine, limb muscle, liver, lung, mammary gland, MAT, pancreas, SCAT, skin, spleen, thymus, tongue, and trachea. AUROC of each cell type is shown in rings.

## Methods

### scRNA-seq datasets

We used the compendium of single cell transcriptomic data from the Tabular Muris Senis^18^. Cell type annotations in Tabula Muris Senis were curated by domain experts and all cell type annotations present in the dataset were manually mapped to the Cell Ontology vocabulary. We next obtained 26 scRNA-seq datasets from 11 different studies^6–10,29–35^. We used the processed collection from Scanorama^26^, where low-quality cells were excluded. There were 5,216 genes across all 26 datasets and a total of 105,476 cells, with each dataset containing between 90 and 18,018 cells. Since these datasets did not provide cell type annotations that were mapped to the Cell Ontology vocabulary, we manually mapped cell types in these datasets to Cell Ontology terms (**Supplementary Table 3**). After the mapping, there were 10 different Cell Ontology terms in these 26 datasets. We denoted these datasets as “26-datasets” in this paper.

### The Cell Ontology

We downloaded the Cell Ontology from The OBO Foundry (http://www.obofoundry.org/ontology/cl.html)^20^. We used the “is_a” relation in the Cell Ontology to construct an undirected graph of cell types. There were in total of 2331 nodes in the constructed graph, corresponding to 2331 different cell types. All edges in this graph have the same weight.

### Embedding the Cell Ontology into the low-dimensional space

OnClass computed a compressed, low-dimensional representation of each cell type based on the constructed cell type graph. We used clusDCA^24,25^, which had been proposed to embed the Gene Ontology, to embed the Cell Ontology. clusDCA first computed a propagated cell type graph by applying the random walk with restart^36,37^ to the cell type graph. It then obtained the low-dimensional representation of each cell type by using the singular value decomposition (SVD)^38^ to reduce the dimensionality of this propagated cell type graph. As suggested by clusDCA, we set the dimensionality of SVD to 1000 and the restart probability of the random walk with restart to 0.8. A detailed description of embedding cell types can be found in the Supplement (**Supplementary Fig. 1, Supplementary Note**).

### Cell type annotation

OnClass used a bilinear neural network model to predict the Cell Ontology term for a novel cell. Let *M* be an *m* by *n* matrix of input gene expression data, where *m* was the number of cells and *n* was the number of genes. Let *Y* be an m by *c* label matrix, where *c* was the total number of Cell Ontology terms in the Cell Ontology. *Y*_*ij*_*=*1 if cell *i* was annotated to Cell Ontology term *j*, otherwise *Y*_*ij*_*=*0. Note that *c* was often much larger than the number of seen Cell Ontology terms in the training data, as the majority of Cell Ontology terms were unseen in the training data. The corresponding columns of unseen Cell Ontology terms were all zeros in the label matrix. Let *X* be a *c* by *q* matrix of the low-dimensional representations of cell types, where *q* was the dimension of cell type embedding dimensionality. *X* was the output of clusDCA and fixed during optimization. OnClass optimized the following cross-entropy loss:

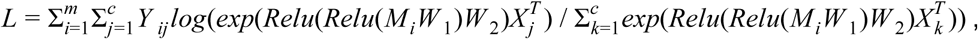

Where *W*_1_ ∈ *R*^*n***×***h*^ and *W*_2_ ∈ *R*^*h***×***q*^ were the parameters that needed to be estimated. *Relu* was the rectifier function for nonlinear transformation^39^. *h* was the number of hidden dimensions and set to 500. We observed that the performance of OnClass was stable for *h* between 200 and 2000. OnClass used ADAM ^40^ to optimize this objective function.

After the optimization, the Cell Ontology term of a new cell with expression vector *z* could then be predicted as:

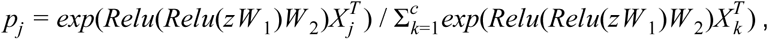

where *p*_*j*_ was the probability that this cell belonged to Cell Ontology term *j. P* = {*p*_1_, *p*_2_, …, *p*_*c*_} was the probability distribution that this cell belonged to each Cell Ontology term, including both seen Cell Ontology terms and unseen Cell Ontology terms. As a result, OnClass could automatically assign cells to any term in the Cell Ontology, even if it does not have any annotated cells in the training data.

### Cell type embeddings reflect cell type similarity

We calculated three types of cell type similarities: the Cell Ontology structure-based similarity, the embedding-based similarity and the gene expression-based similarity. The Cell Ontology structure-based similarity was calculated as the shortest distance between two cell types in the Cell Ontology graph. The embedding-based similarity was the cosine similarity between low-dimensional representations of two cell types. We used the gene expression of all FACS cells in TMS to calculate the gene expression-based similarity. The calculation was performed per organ. For each organ, we first identified two sets of cells belonging to two given cell types. We then calculated the mean of pairwise cosine similarities between gene expression of these two sets of cells and used it as the gene expression-based cell type similarity.

### Evaluation of cell type annotation

We evaluated across different proportions of seen Cell Ontology terms in the test set ranging from 100% to 10%, where 10% indicates that 10% of Cell Ontology terms in the test set have at least some annotated cells in the training data. For a proportion *k* percentage, we first randomly selected *k* percentage of Cell Ontology terms as seen Cell Ontology terms and remaining Cell Ontology terms as unseen Cell Ontology terms. All cells belonging to these unseen Cell Ontology terms were used as the test set. For the seen cell types, we random split their cells into five equal size folds, where one-fold was used as the training set and the remaining four-folds were used as the test set. We created a five-fold of test and training here according to the initial annotation process in Tabula Muris Senis, where about 20% of cells (3-month mice) were annotated first and then extended to the remaining 80%. The test data thus contained all cells in each of the unseen Cell Ontology terms and 80% of cells in each seen Cell Ontology terms. We performed cross-validation by repeating this procedure 5 times for each proportion.

To evaluate the case where all Cell Ontology terms in the test set are unseen (**Fig. 3a**), we compared the performance across different proportions of seen Cell Ontology terms in the training set. For a given proportion *k* percentage, we randomly selected *k* percentage of cell types as seen Cell Ontology terms and the remaining as unseen Cell Ontology terms. All cells belonging to the seen (unseen) Cell Ontology terms were used as the training (test) set. We performed cross-validation by repeating this procedure 5 times for each proportion.

We evaluated our method and comparison approaches on four metrics, including the area under the receiver operating characteristic curve (AUROC), Accuracy@3, Accuracy@5, and Cohen’s kappa statistic^41^. As we were evaluating a large number of classes (i.e., more than 80 cell types), it was important to address the bias from class imbalance during evaluation. Therefore, we used the macro-average AUROC rather than the micro-average AUROC to summarize results across different Cell Ontology terms. Macro-average AUROC calculates the areas under the curves for each class independently and then takes the average. Cohen’s kappa statistic can handle well both multi-class and imbalanced class problems and has been widely used as an alternative to accuracy^11^. A large cohen’s kappa statistic indicates better performance, while 1 indicates perfect classification. Accuracy@3 (Accuracy@5) is a widely used ranking metric, which assesses the correctness of the top 3(5) predicted Cell Ontology terms in comparison to only examining the top 1 Cell Ontology term in Cohen’s kappa statistic. A prediction would be deemed as correct if any of the top 3 (5 for Accuracy@5) predicted Cell Ontology terms is the correct Cell Ontology term.

### Comparison approaches

We compared our method with four existing methods ACTINN, singleCellNet (sCN), one-vs-rest logistic regression (LR), and DOC. ACTINN and sCN are two of the best approaches in cell type annotation according to a recent study^15^. ACTINN used a three-layer neural network to predict the cell type^13^. We used the implementation of ACTINN from the authors (https://github.com/mafeiyang/ACTINN) and ran it on TMS. We used the default parameters for ACTINN since these parameters were used in their paper to annotate cells in the Tabula Muris^3^, an earlier version and subset of our dataset. sCN used gene pairs as features and random forest as the classifier to predict the cell type^11^. Notably, sCN was able to classify cells into an unknown cell type. We obtained the implementation of singleCellNet from (https://github.com/pcahan1/singleCellNet). We found that the implementation of sCN was not scaled to large datasets like TMS and it was not able to cross-validate rare cell types with less than 50 cells. We reimplemented part of sCN to enable its annotation for rare cell types and made the code available as part of our package. To make it scalable to TMS, we ran it on the dimensionality reduced gene expression matrix instead of the original gene expression matrix. LR was the standard machine learning classifier for multi-class classification on large-scale datasets. We used the one-vs-rest logistic regression instead of the multinomial logistic regression in order to obtain a probability cutoff of 0.5 to determine the unknown cell type. DOC was an advanced machine learning method for classifying unseen text documents, which was a natural solution to our problem and could be directly applied here^42^. The key idea of DOC was to find a data-driven probability cutoff for the unknown class rather than using a fixed probability cutoff of 0.5 as LR did. However, DOC was also not able to classify cells into the specific cell type. As the original DOC codebase was developed for word sequences classification and could not directly take gene expression as input, we reimplemented and replaced its underlying convolutional neural network classifier with a multinomial logistic regression.

Although sCN, DOC and LR were able to classify cells into a “unknown” cell type, they were not able to classify these cells into the specific cell type. To enable a fair comparison, we further proposed to extend these three approaches by classifying cells belong to the unknown cell type to a specific cell type. In particular, when a cell was annotated as the unknown cell type, we first found the seen cell type that had the largest confidence score for this cell. We then annotated the cell to the nearest unseen cell type of this seen cell type based on the Cell Ontology graph. We denoted these extended approaches as sCN (extended), LR (extended), and DOC (extended) for sCN, LR, and DOC, respectively.

### Transfer annotations to 26-datasets

To transfer annotations from TMS to 26-datasets, we first used Scanorama to correct batch effects among TMS and 26 datasets. Scanorama took the gene expression matrix of these 27 datasets as input, it then calculated the corrected gene expression of these 27 datasets. We then ran OnClass on all cells in TMS and predicted the Cell Ontology term for each cell in the 26-datasets. To combine these 26-datasets, we used the output probability distribution of each cell by OnClass as the feature for each cell. We visualized these cells by projecting these features using UMAP^43^. We used silhouette coefficients to evaluate the clustering accuracy for both our method and Scanorama^27^.

### Marker genes identification

We used differential gene expression analysis to identify marker genes for each Cell Ontology term. In particular, we first ran OnClass on all FACS cells in TMS and then predicted the probability of these cells belonging to each Cell Ontology term in the Cell Ontology. For each Cell Ontology term, we took the 50 cells with the highest probability as the positively annotated group and the 50 cells with the lowest probability as the negatively annotated group. We then used the t-test to test whether an individual gene was significantly overexpressed in the positively annotated group then the negatively annotated group. We performed this one-sided independent t-test for each gene and then ranked genes according to the resulted *P*-values. This rank list was the predicted marker gene list. Curated marker genes of 69 Cell Ontology terms were collected from literature by experts (**Supplementary Table 4**). 28 cell types in TMS are in these 69 Cell Ontology terms and thus had curated marker genes. To classify a new cell according to marker genes, we used the sum of the expression of marker genes of each Cell Ontology term as the predicted score for that Cell Ontology term. A larger score indicated that the cell more likely belonged to this Cell Ontology term.

### Statistical analysis

We used the scipy.stats^44^ Python package implementation of the one-sided independent t-test, Pearson correlation statistics, Spearman correlation statistics, and associated P-values used in this study. We used the scikit-learn Python package implementation of one-vs-rest logistic regression, silhouette coefficients, AUROC, and cohen’s kappa statistics used in this study^45^.

## Data availability

All datasets used in this study are available at https://figshare.com/projects/OnClass/70637, including gene expression data, pre-trained model, cell type embeddings, and the Cell Ontology. A detailed description of these datasets can be found at https://onclass.readthedocs.io/.

## Code availability

OnClass codes are available at https://github.com/wangshenguiuc/OnClass. The OnClass server can be found at http://onclass.ds.czbiohub.org/.

## Acknowledgments

The authors would like to thank the developers and maintainers of the Cell Ontology for insightful discussions. This work is supported by the Chan-Zuckerberg Biohub, NIH GM102365, LM005652, and TR002515.

