## Supplementary figures for "Unifying single-cell annotations based on the Cell Ontology"

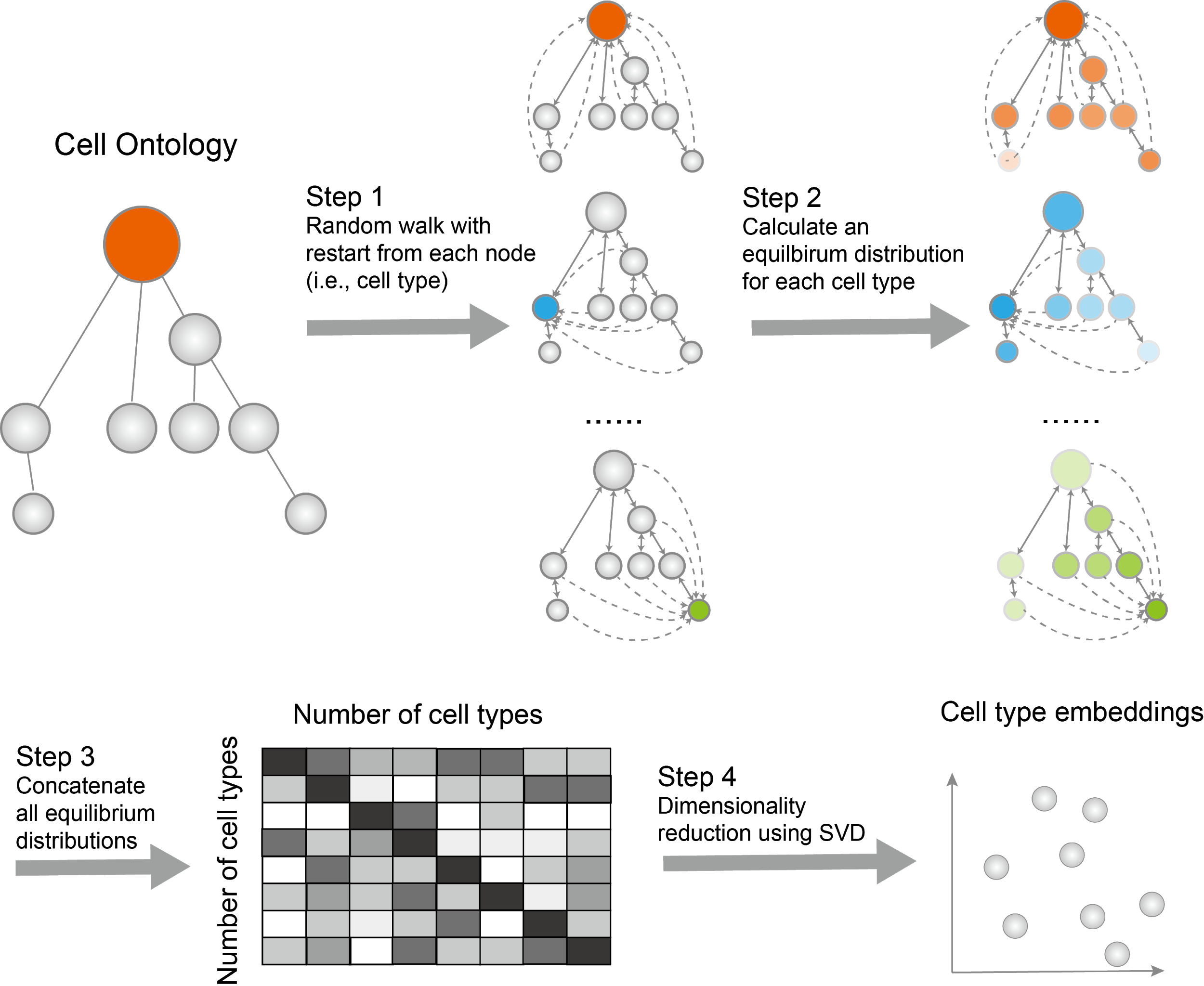

### **Supplementary Figure 1**

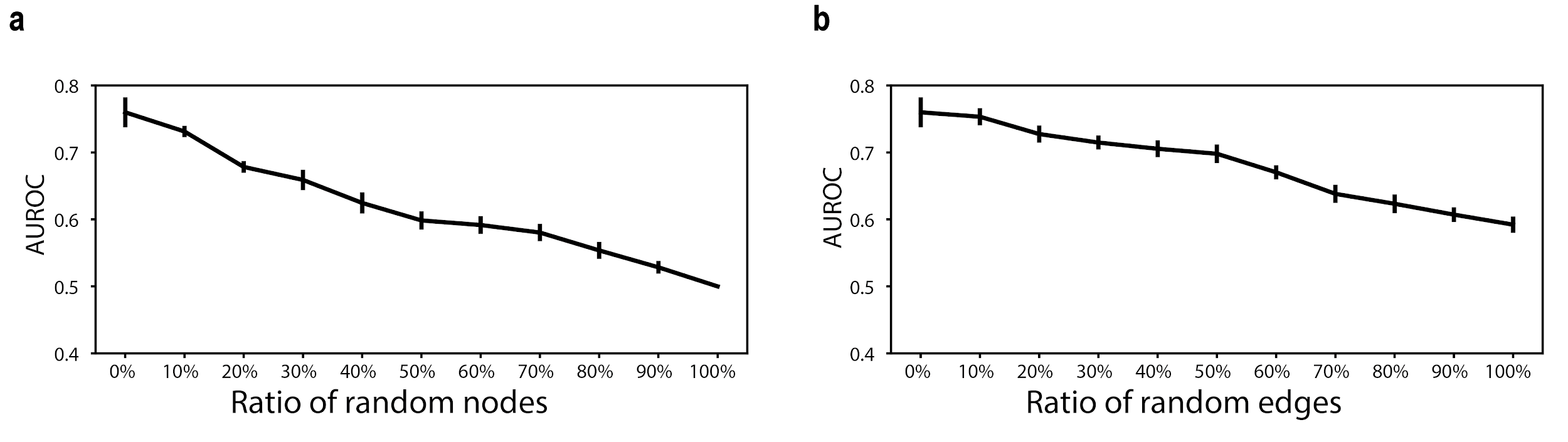

### **Supplementary Figure 2**

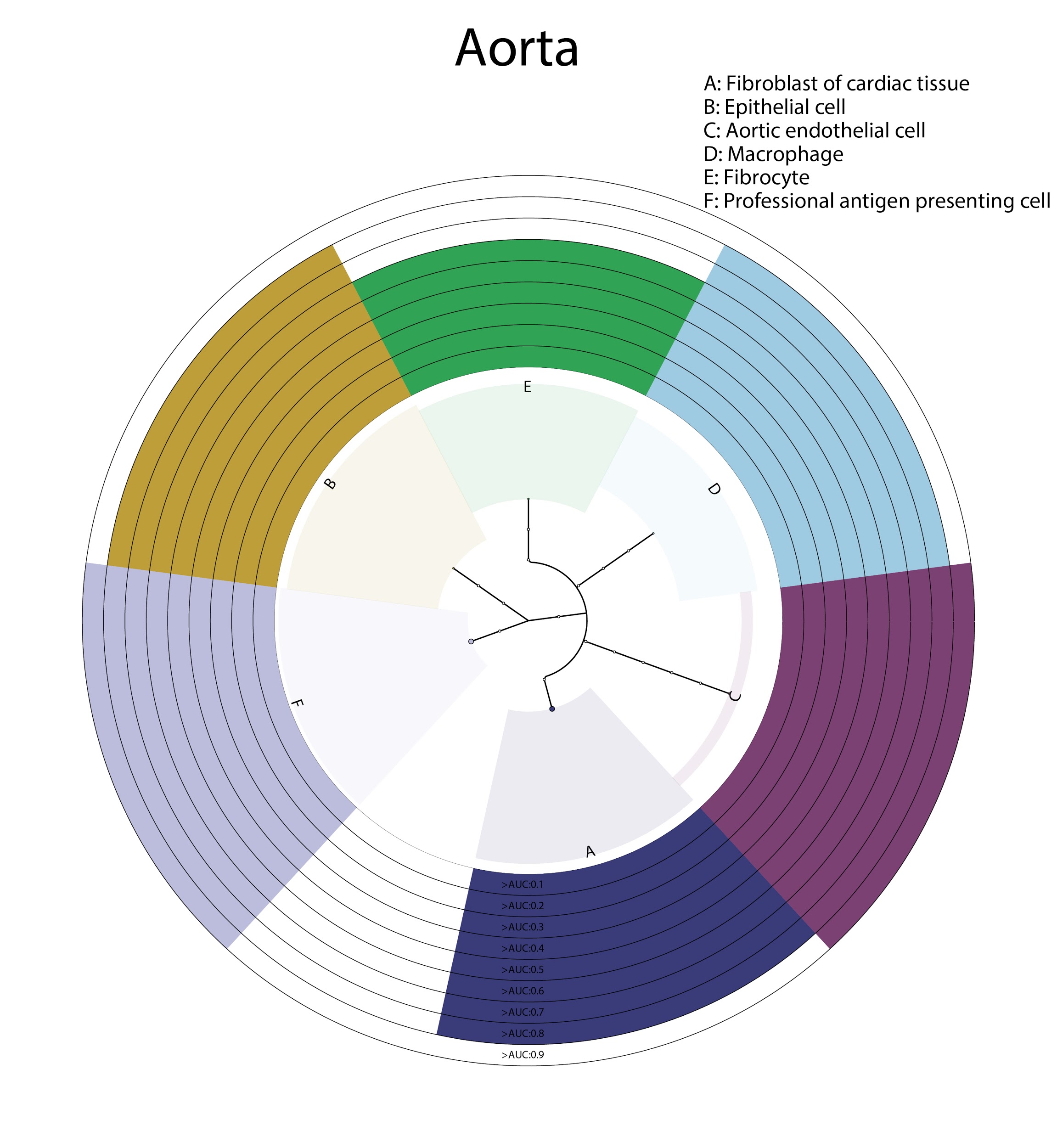

### **Supplementary Figure 3**

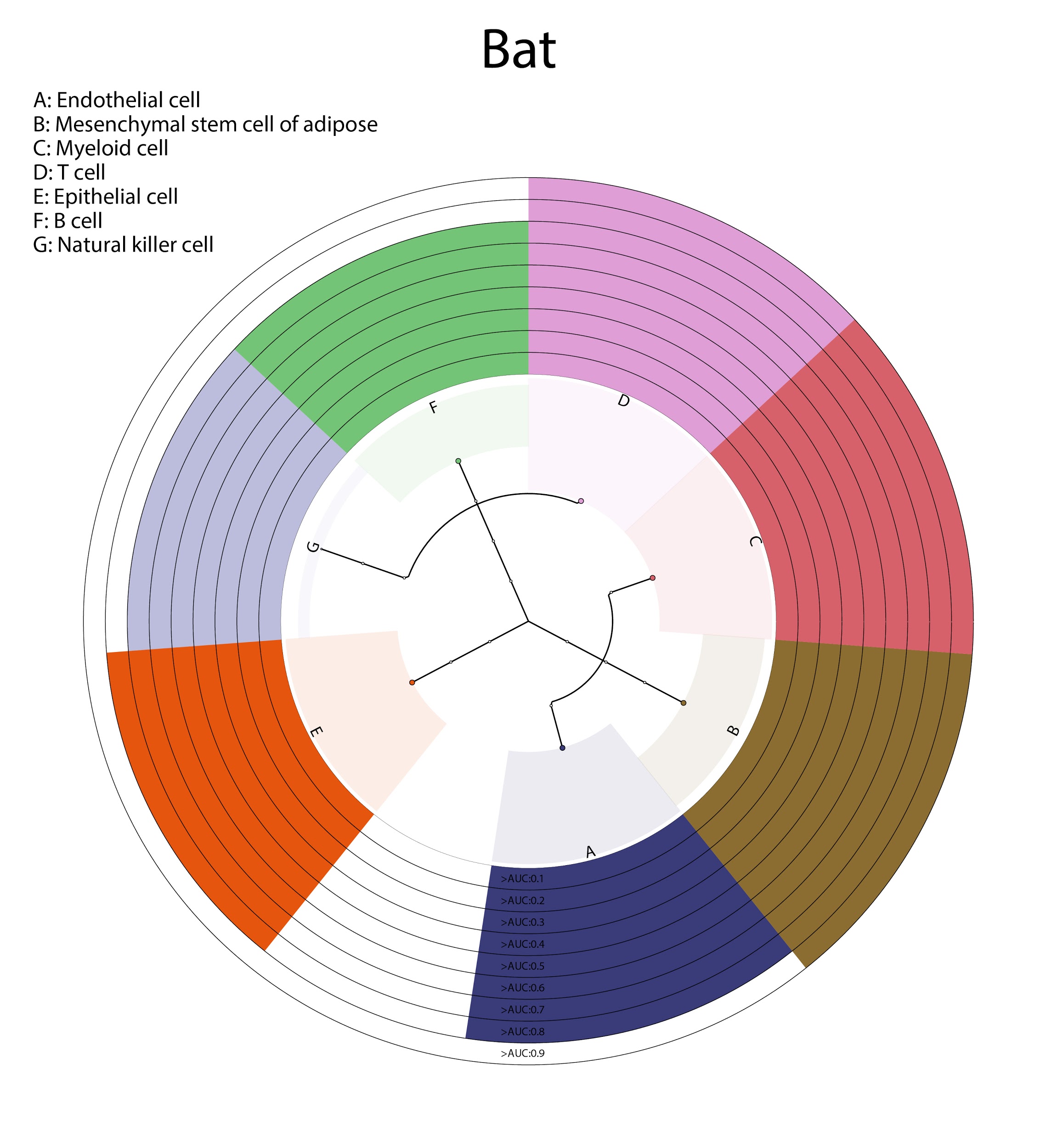

### **Supplementary Figure 4**

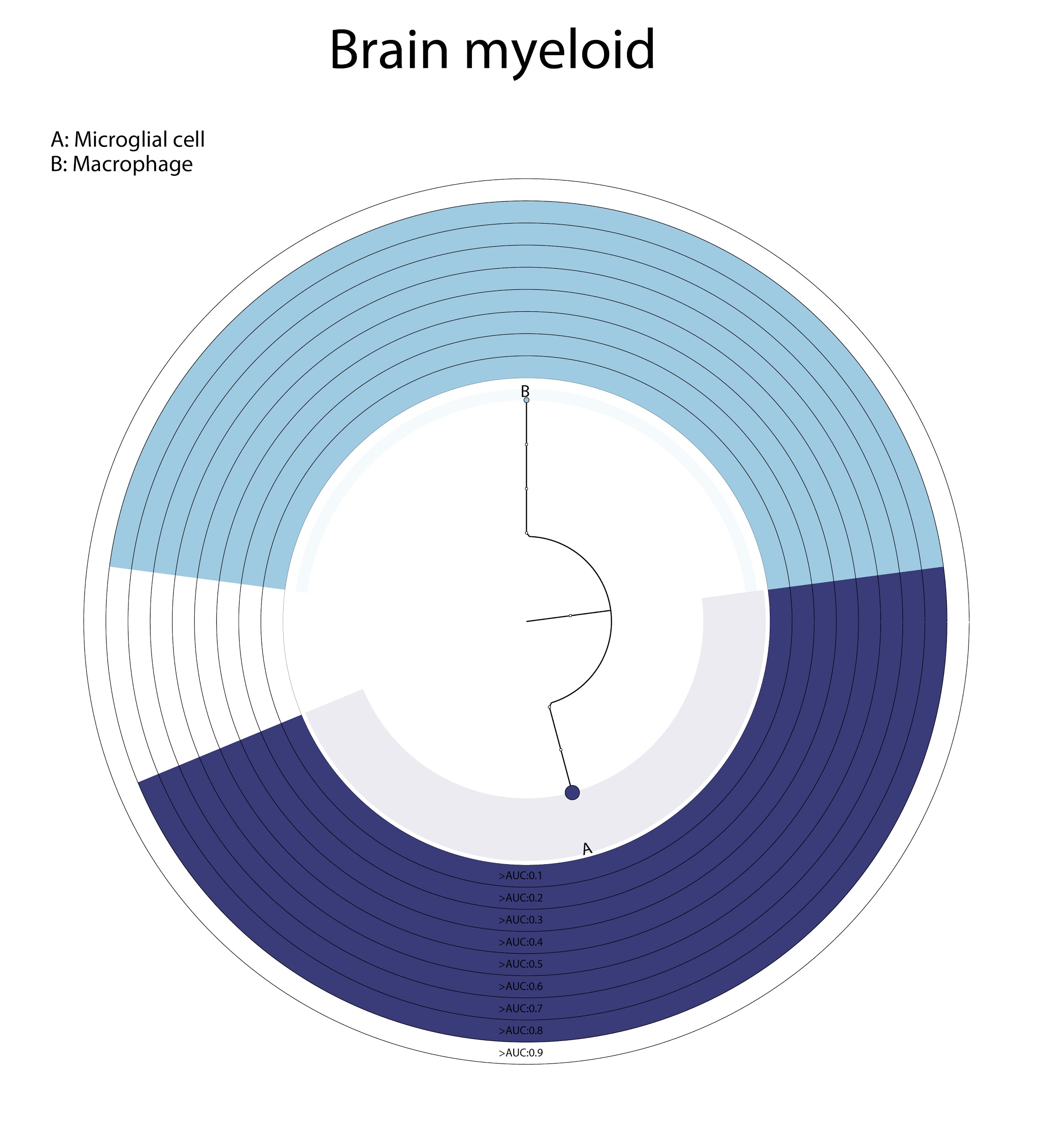

### **Supplementary Figure 5**

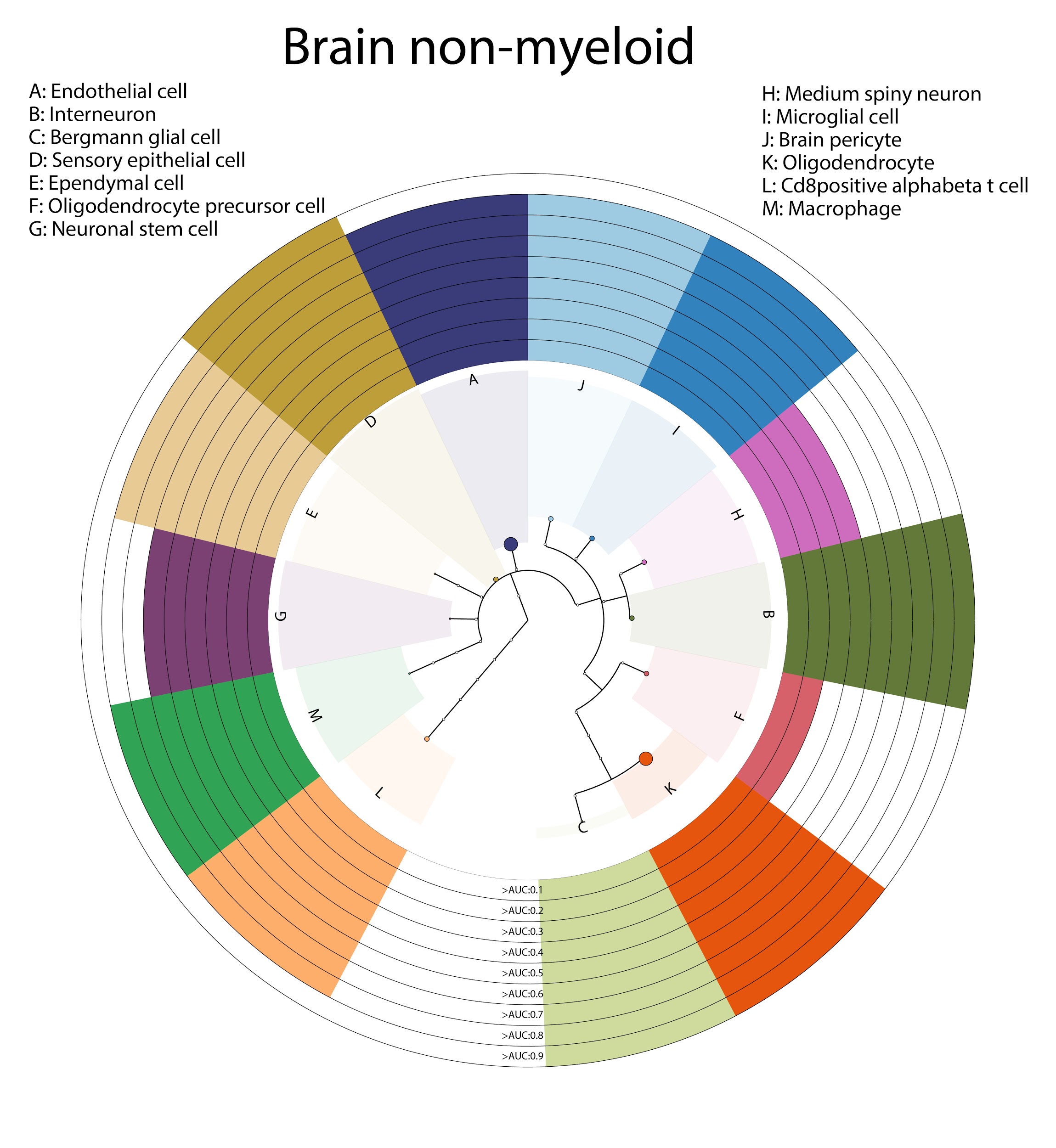

### **Supplementary Figure 6**

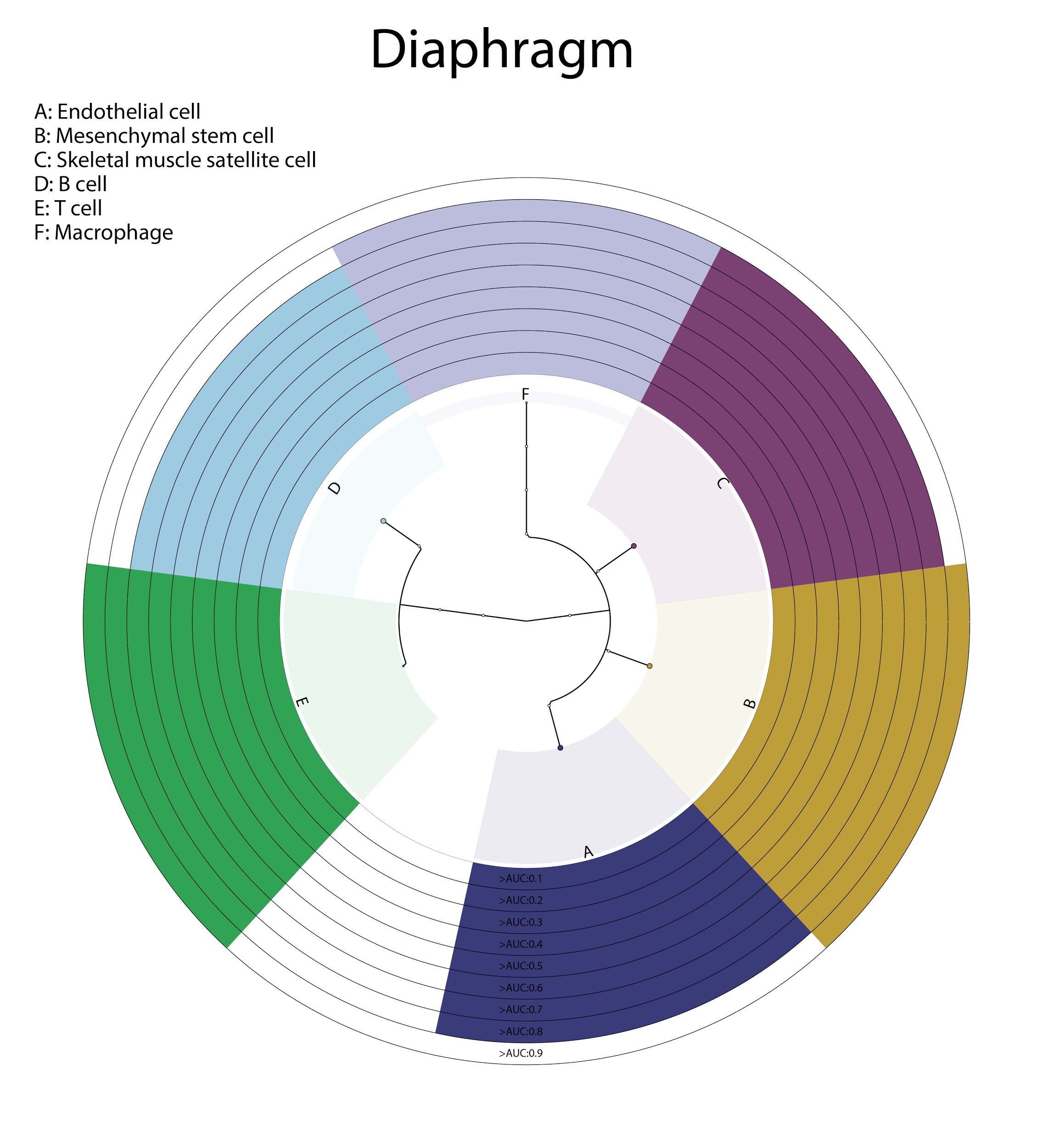

### **Supplementary Figure 7**

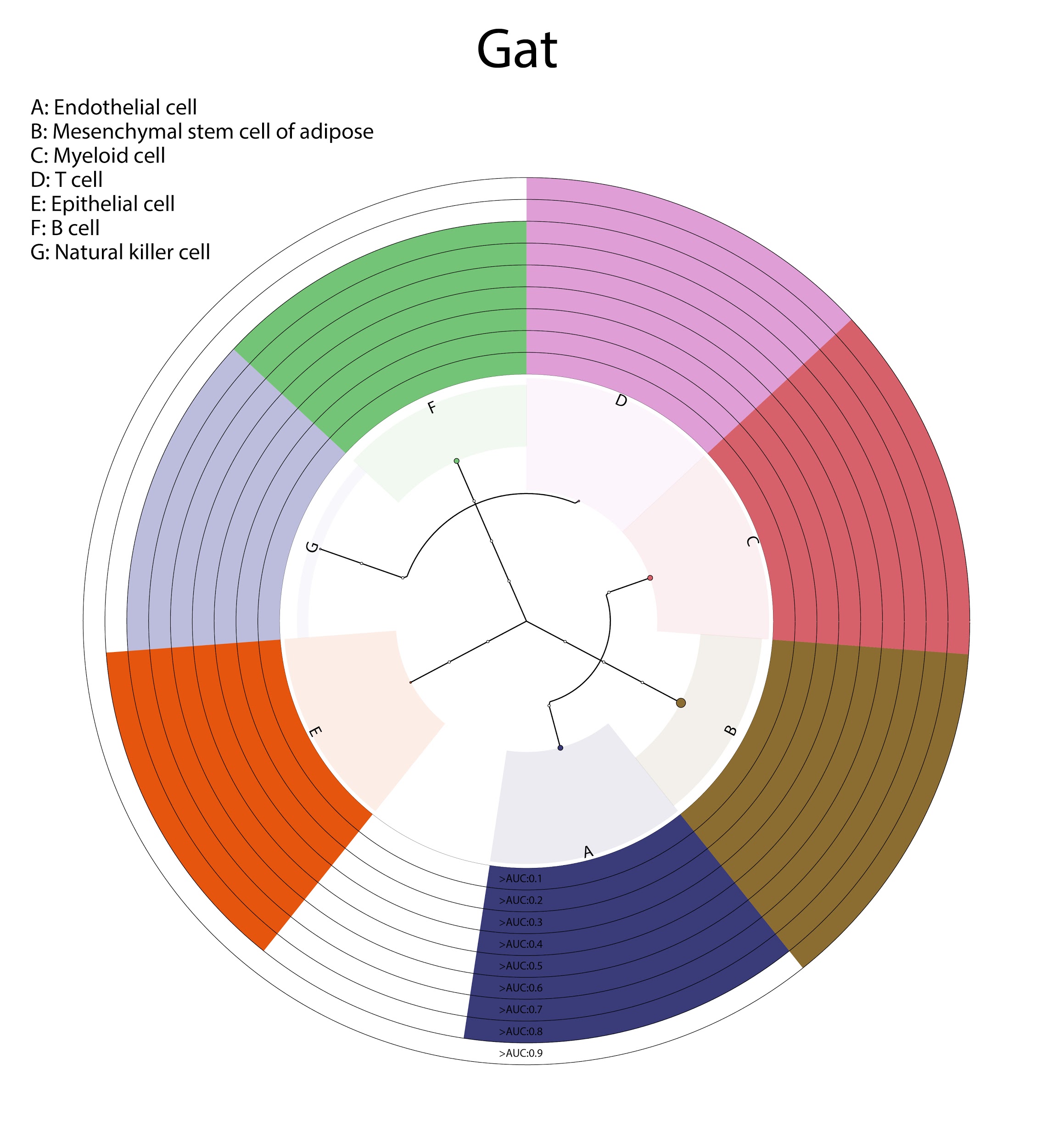

### **Supplementary Figure 8**

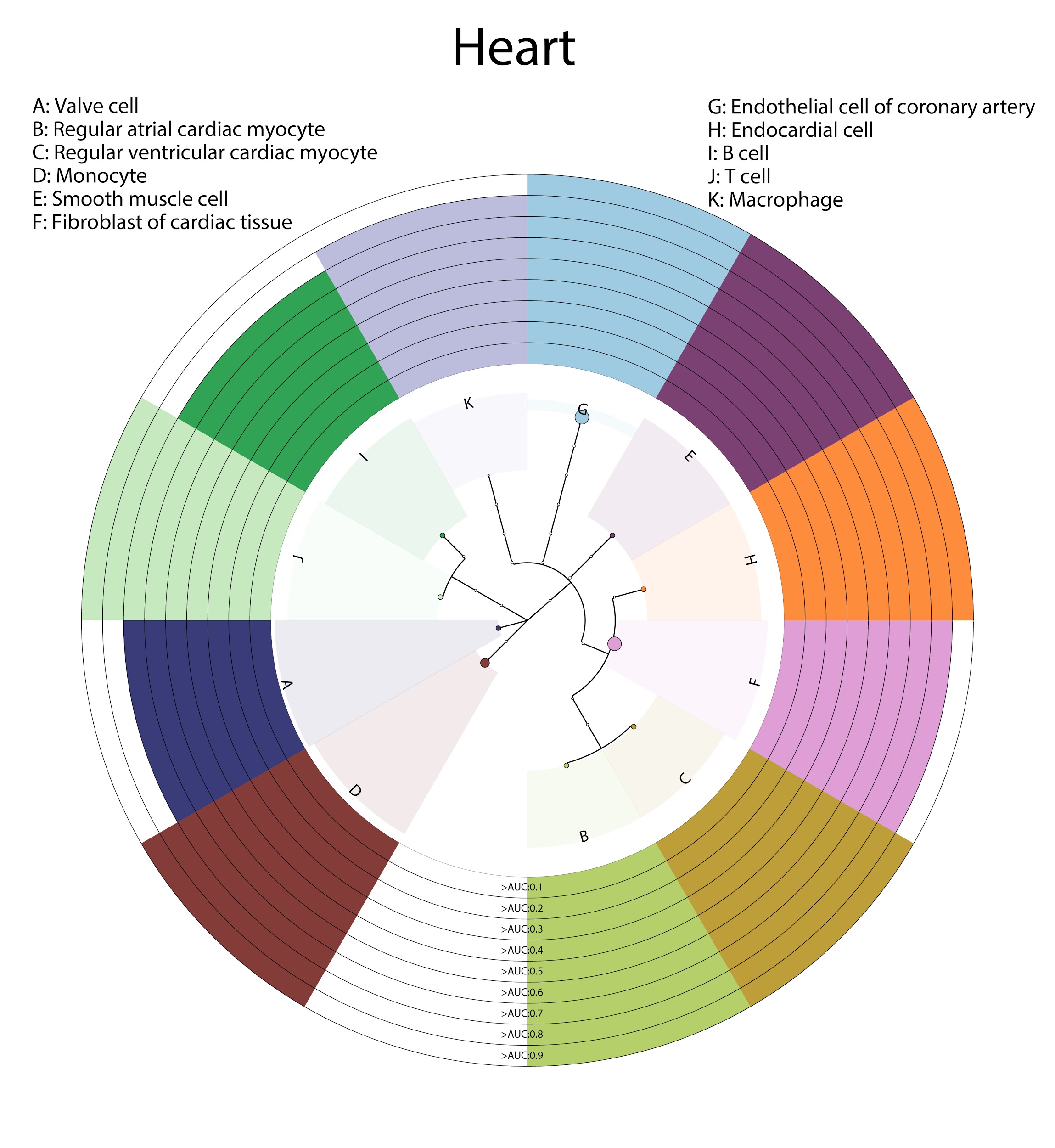

### **Supplementary Figure 9**

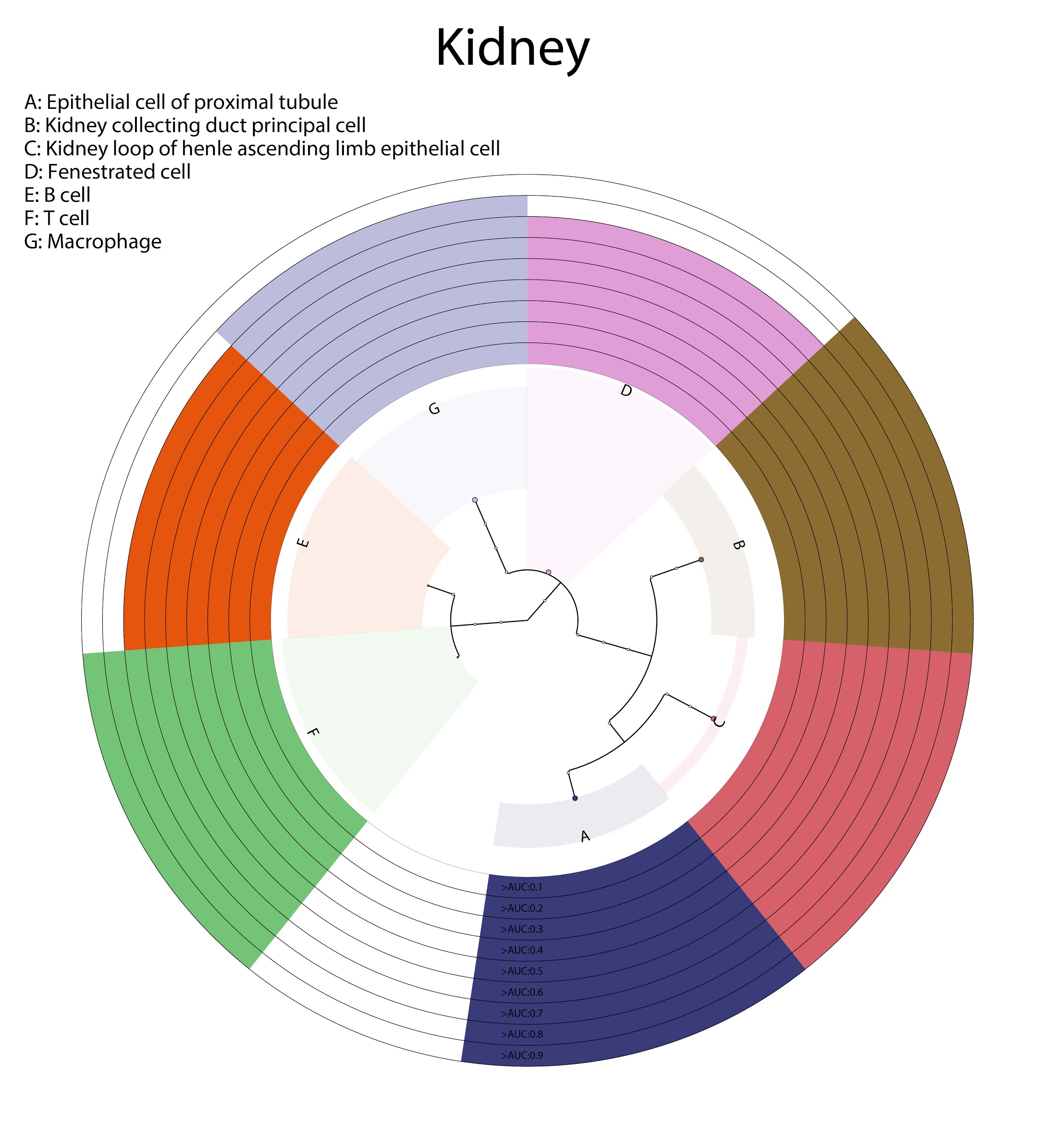

### **Supplementary Figure 10**

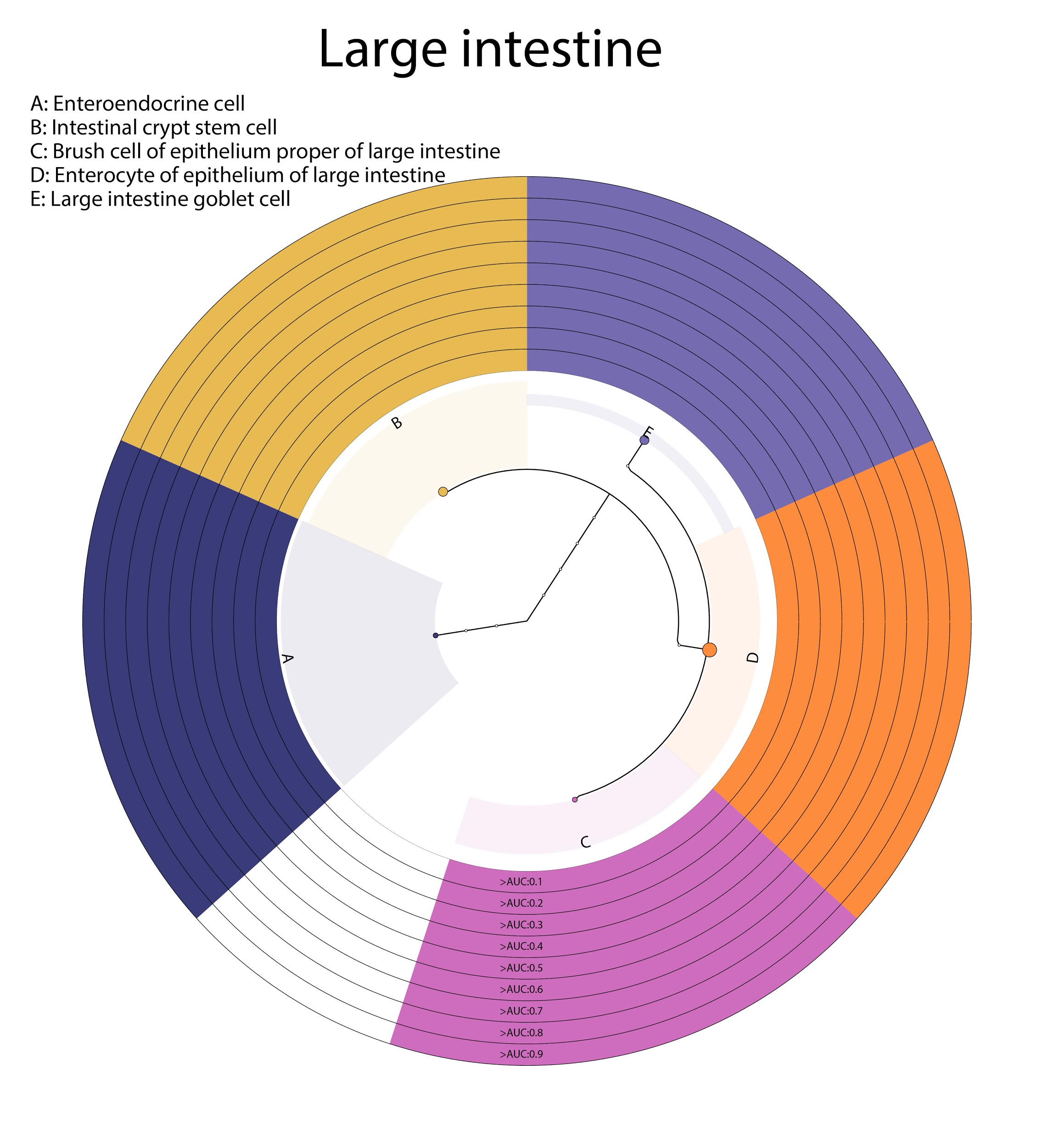

### **Supplementary Figure 11**

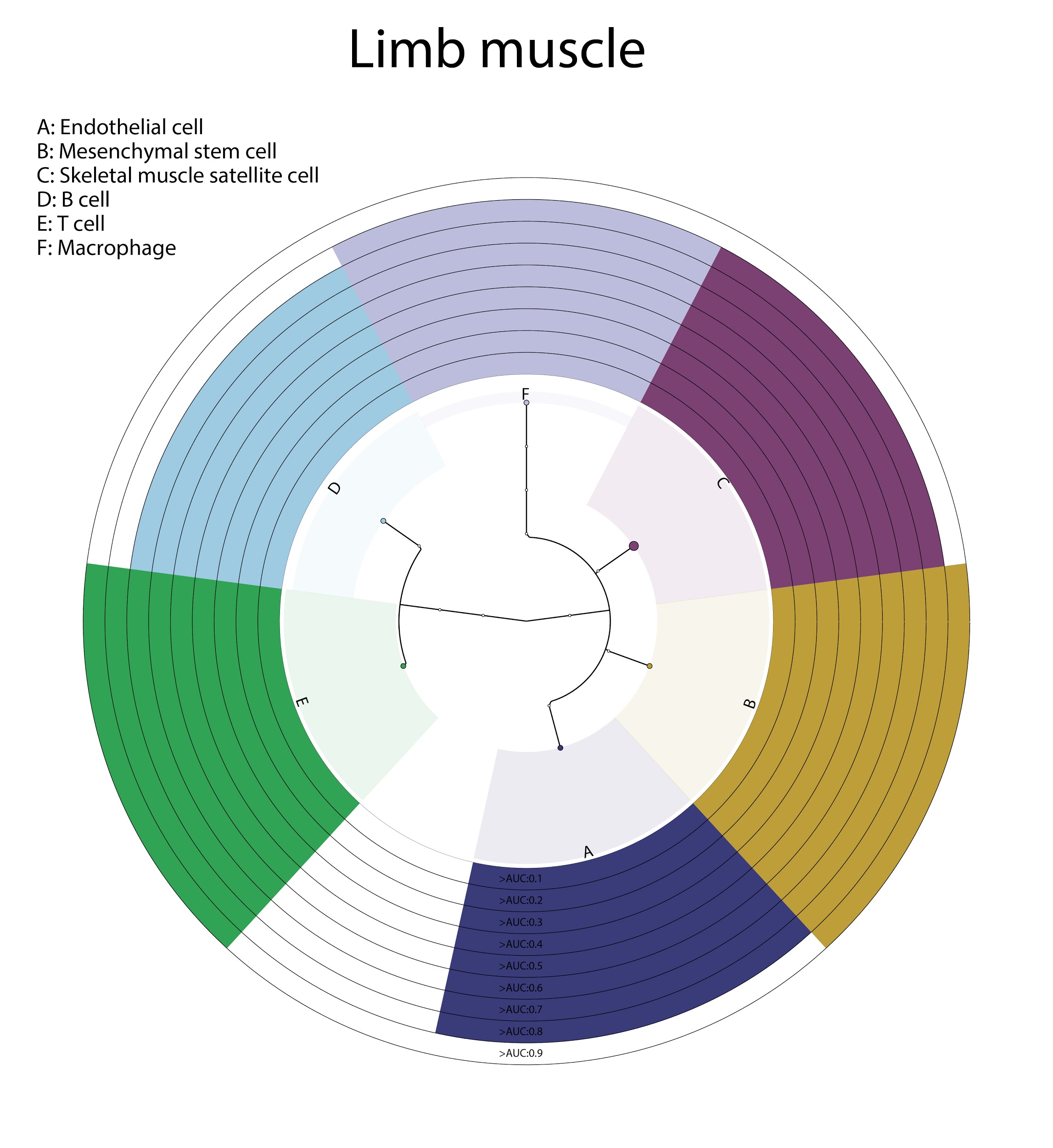

### **Supplementary Figure 12**

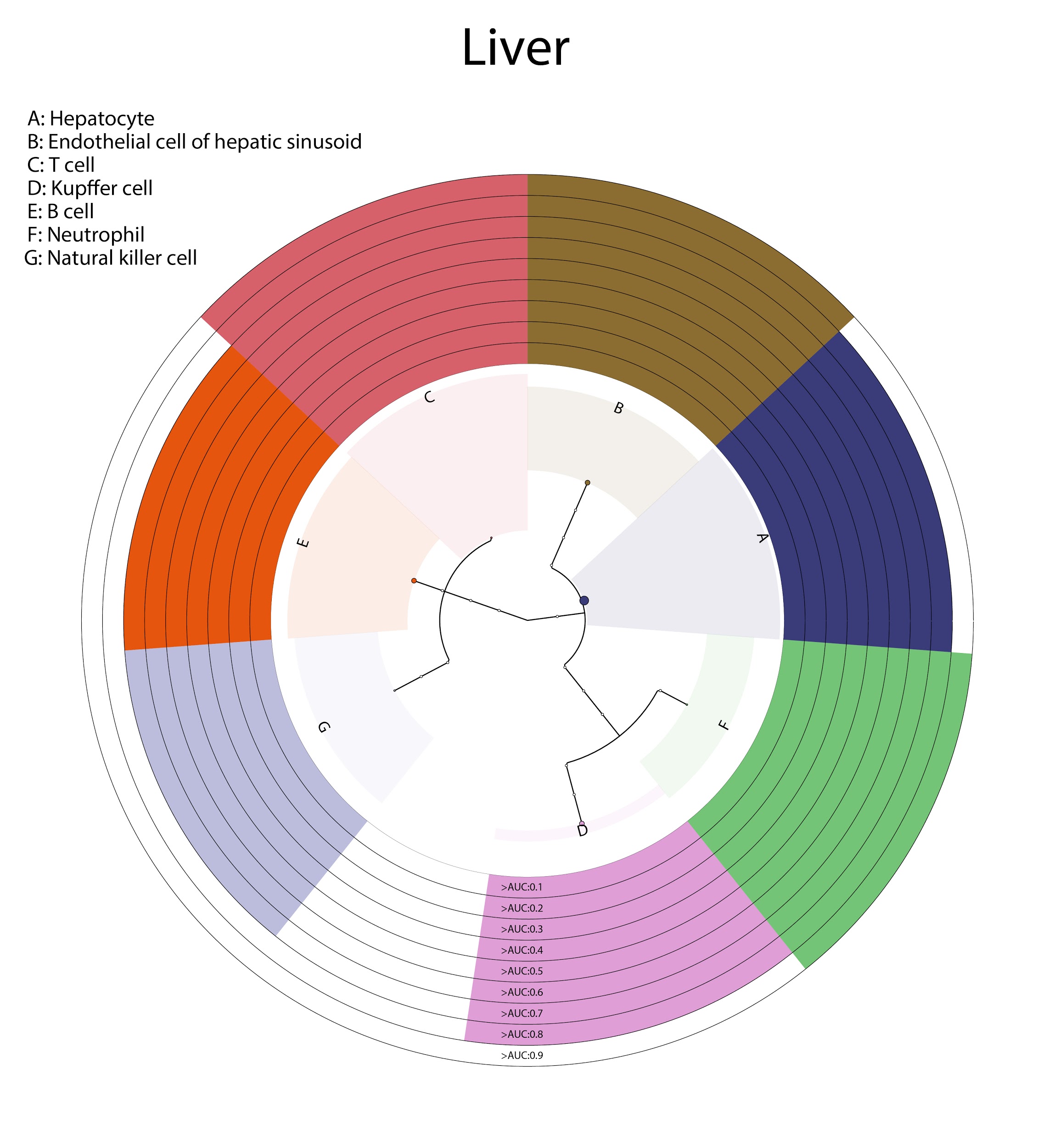

### **Supplementary Figure 13**

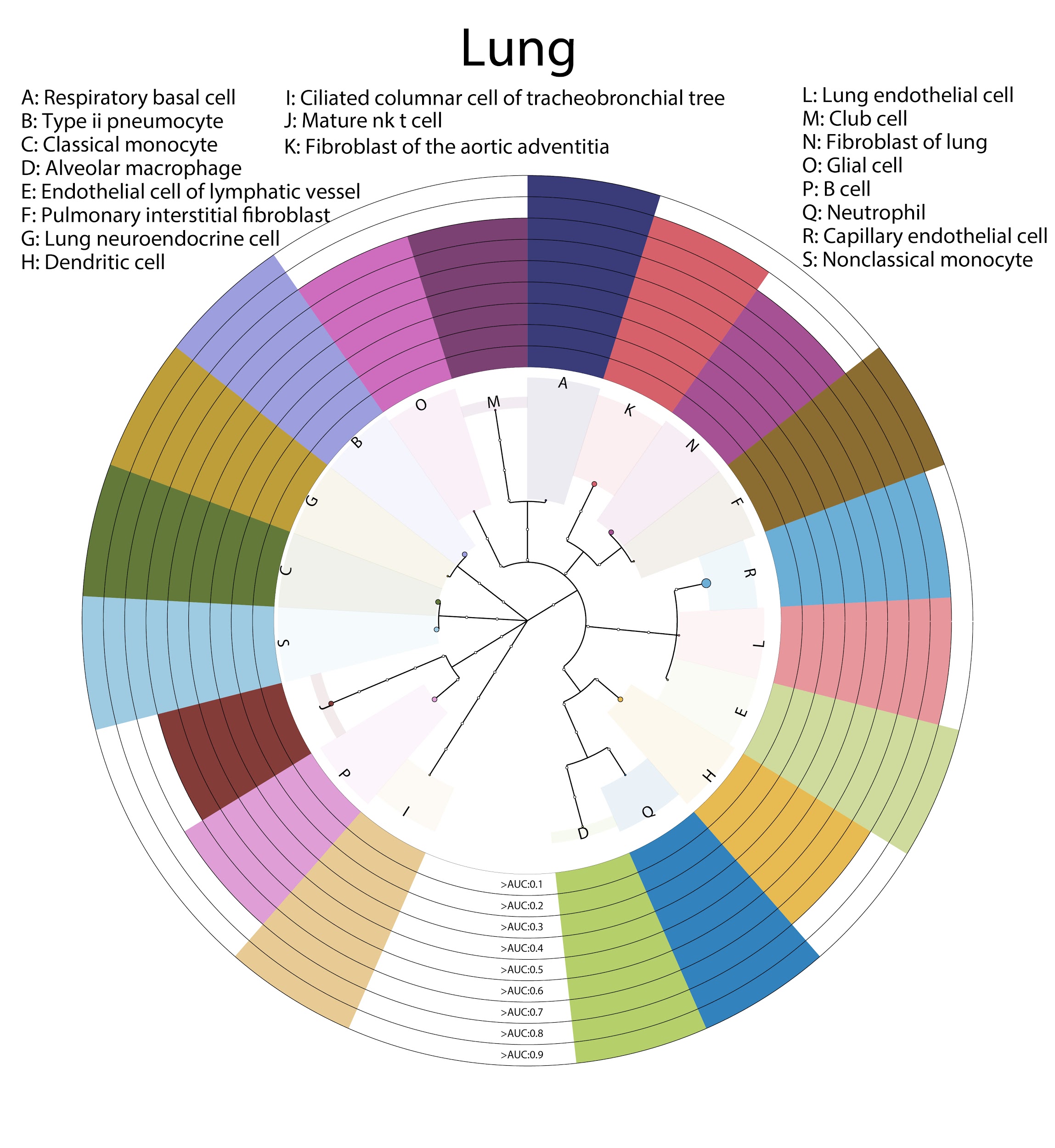

### **Supplementary Figure 14**

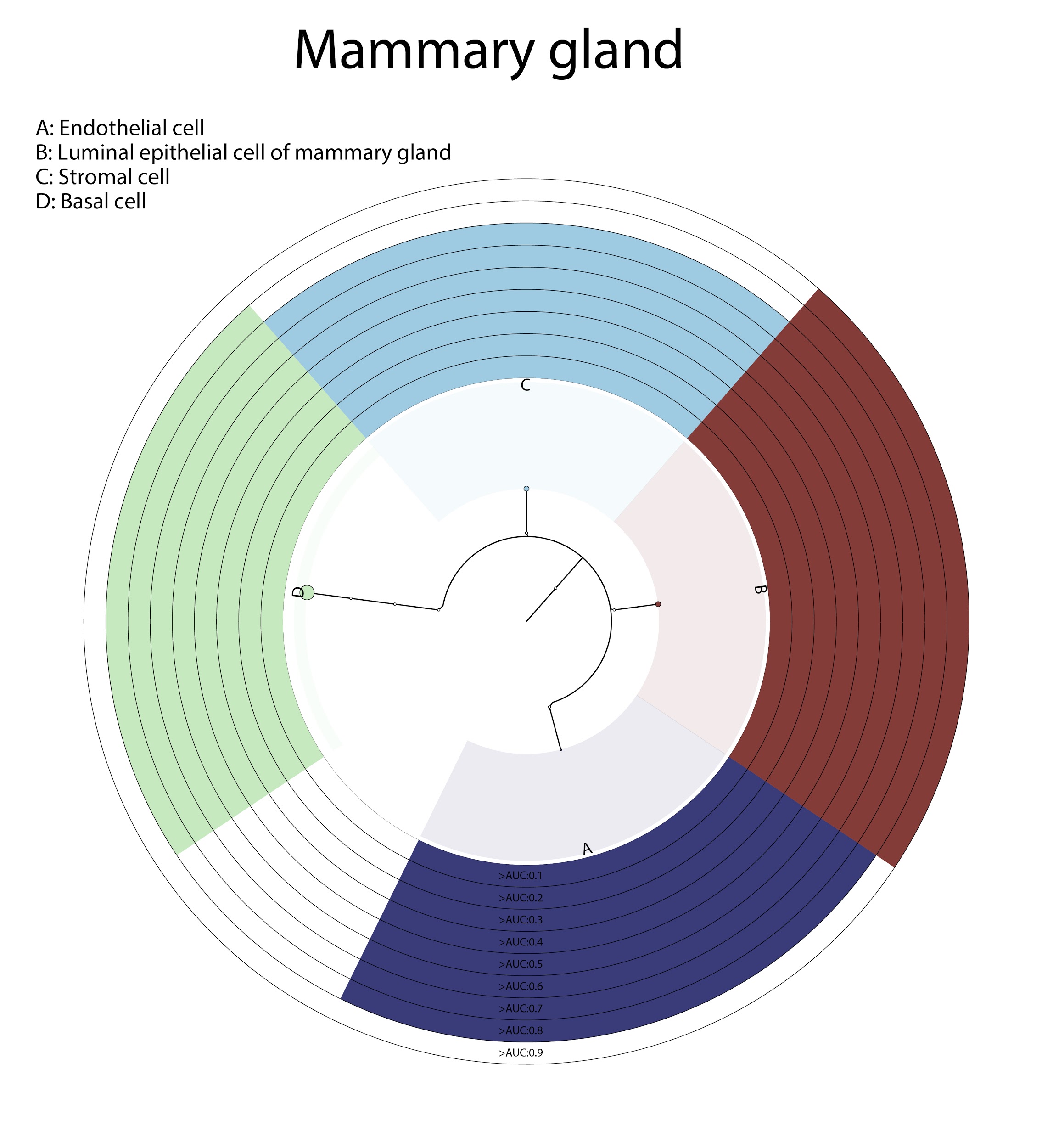

### **Supplementary Figure 15**

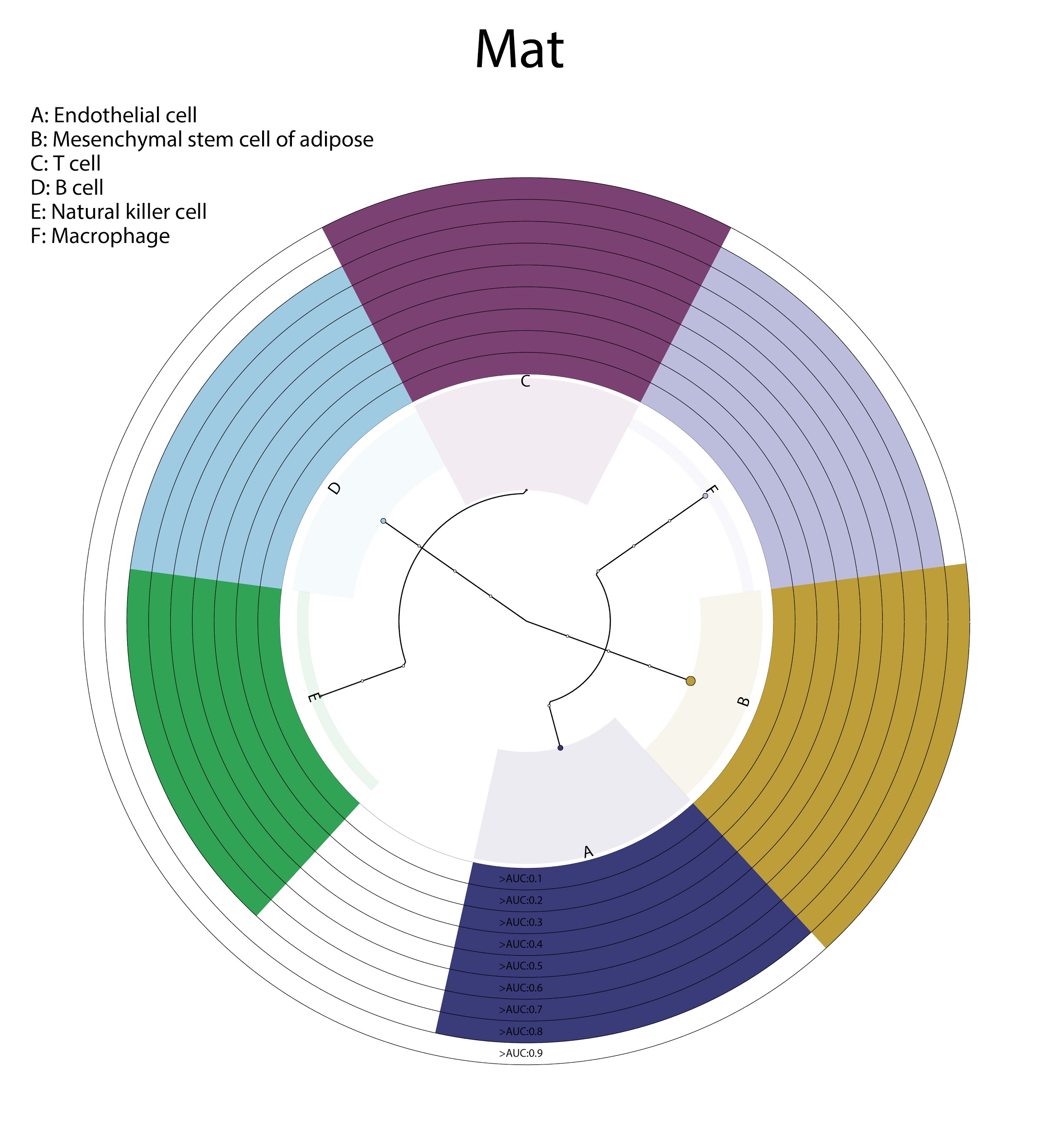

### **Supplementary Figure 16**

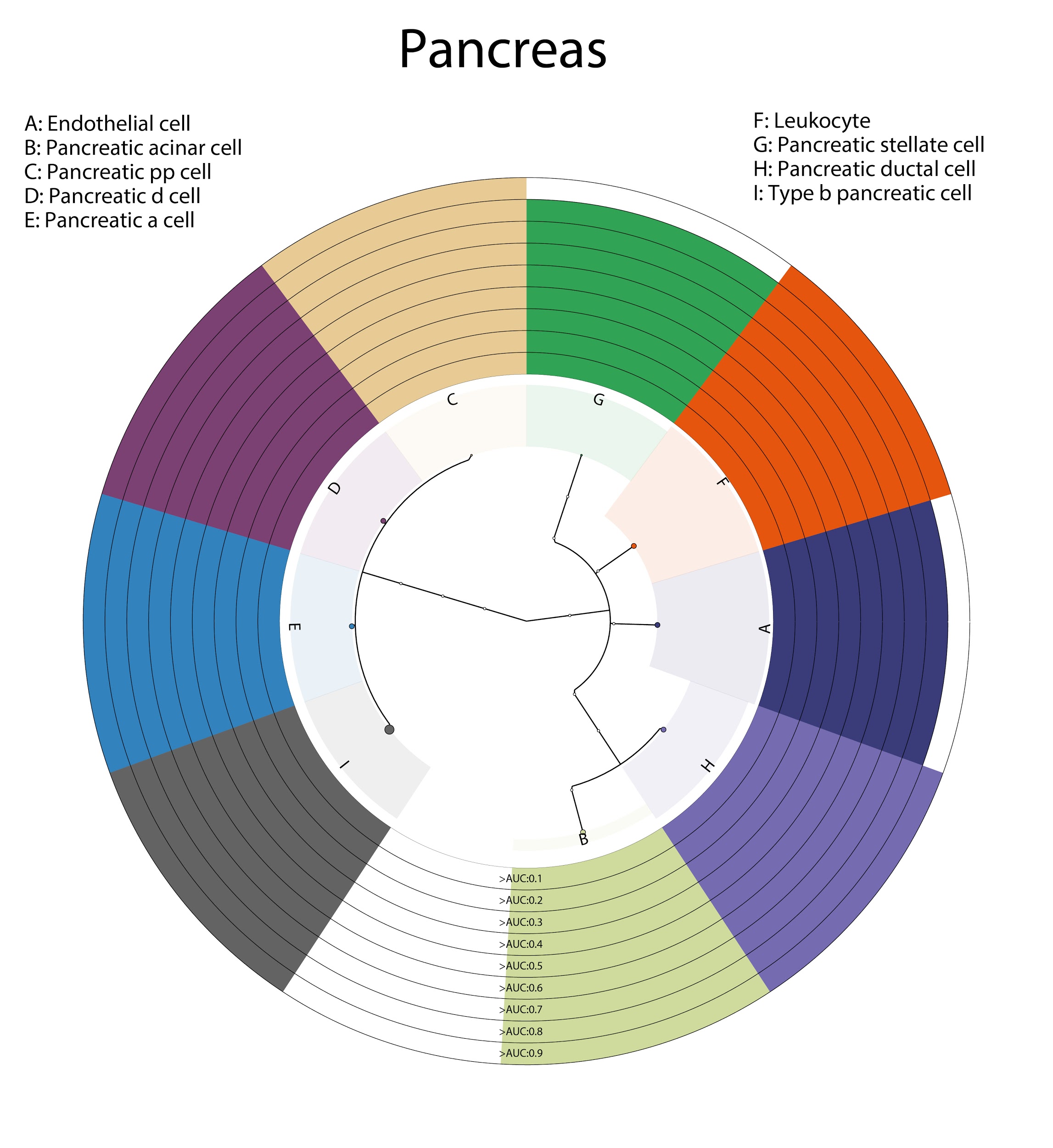

### **Supplementary Figure 17**

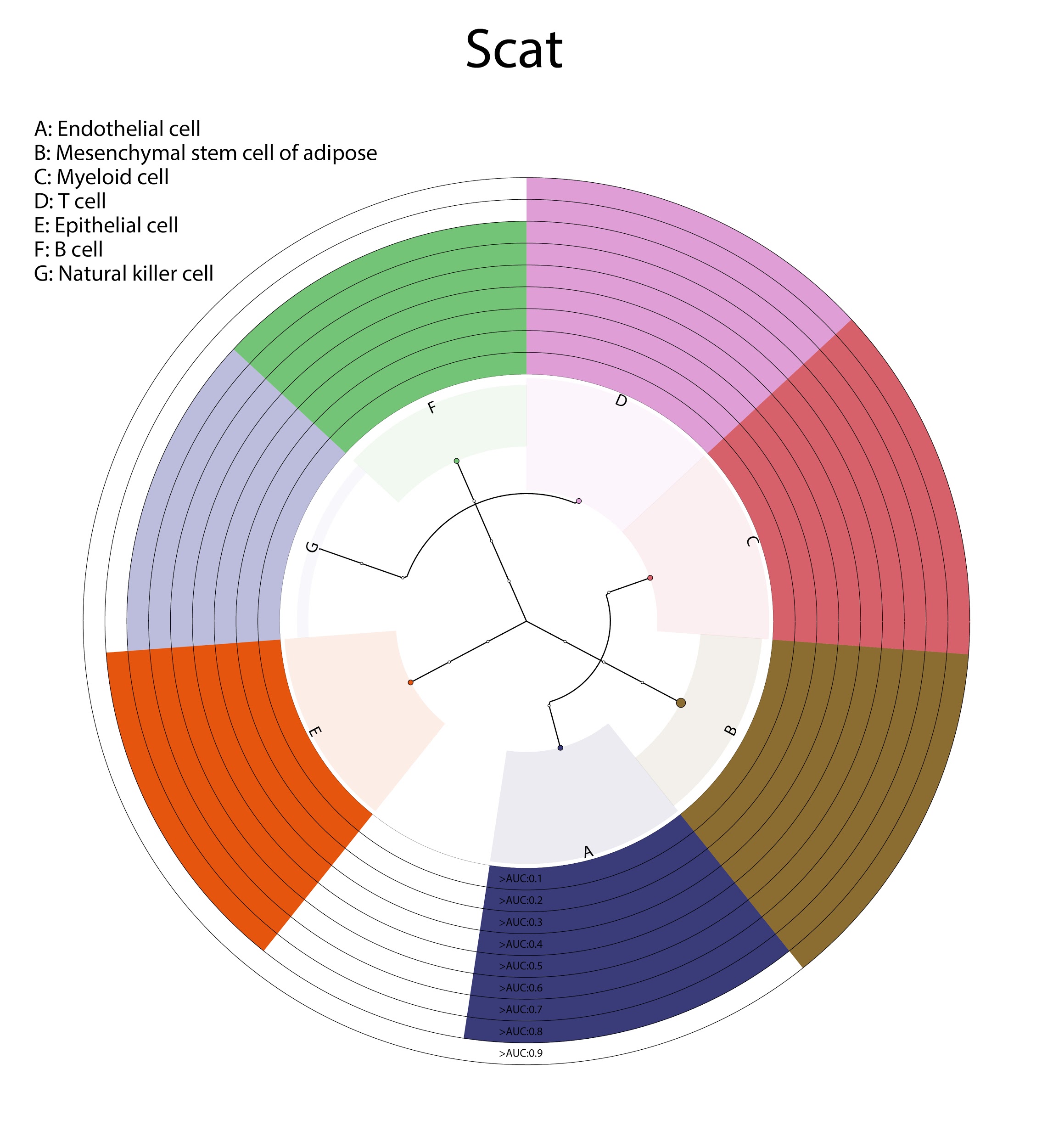

### **Supplementary Figure 18**

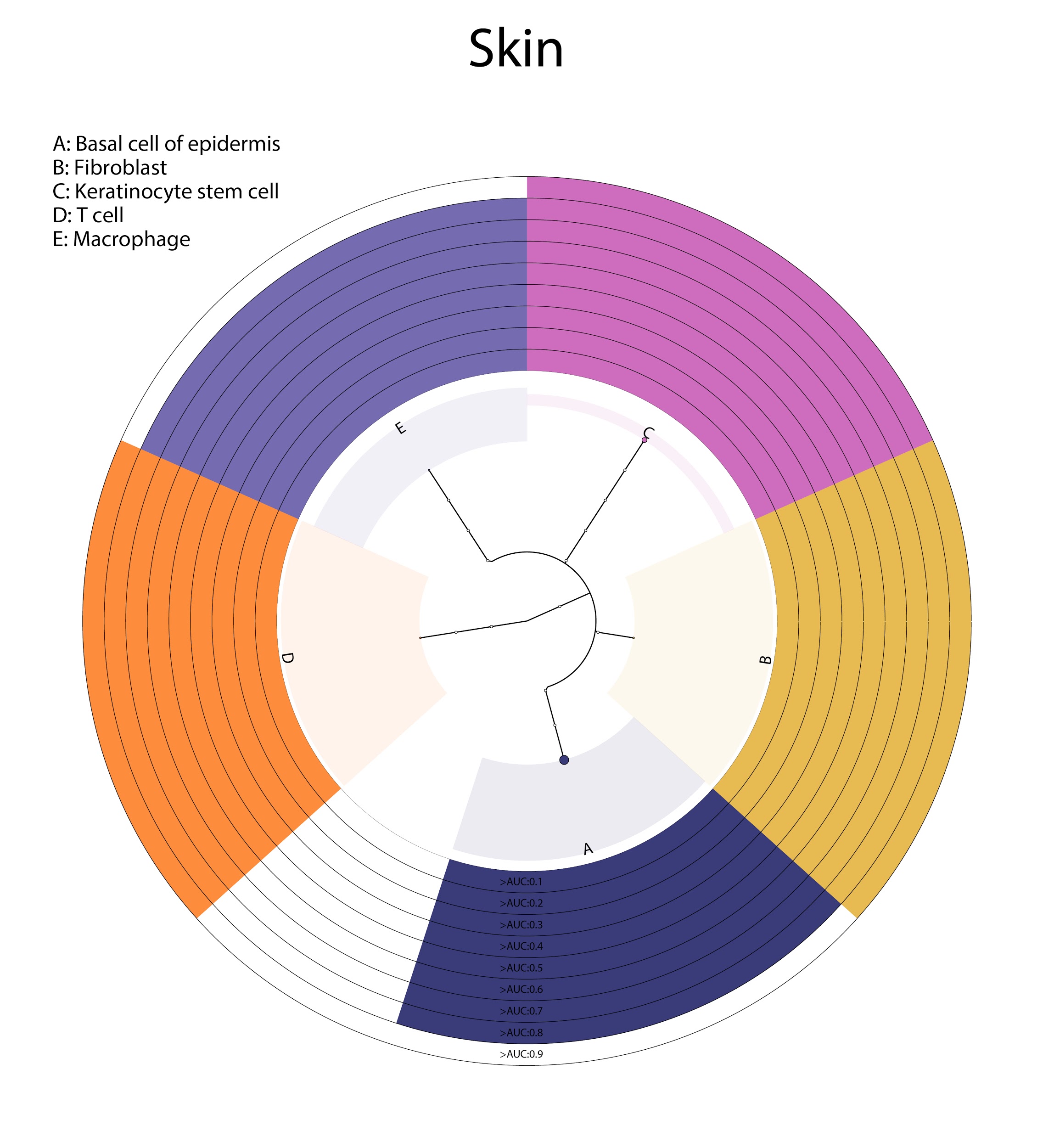

### **Supplementary Figure 19**

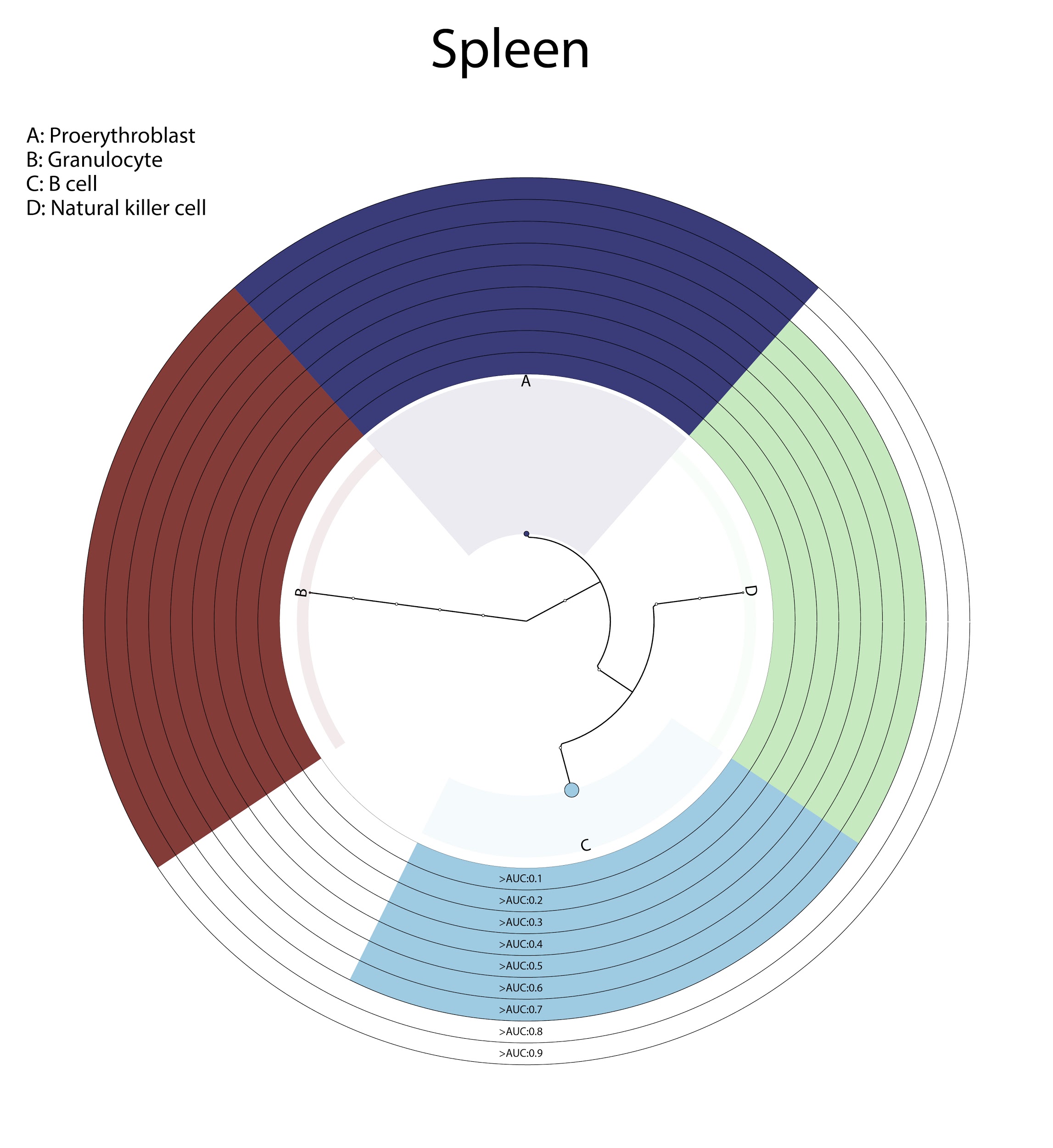

### **Supplementary Figure 20**

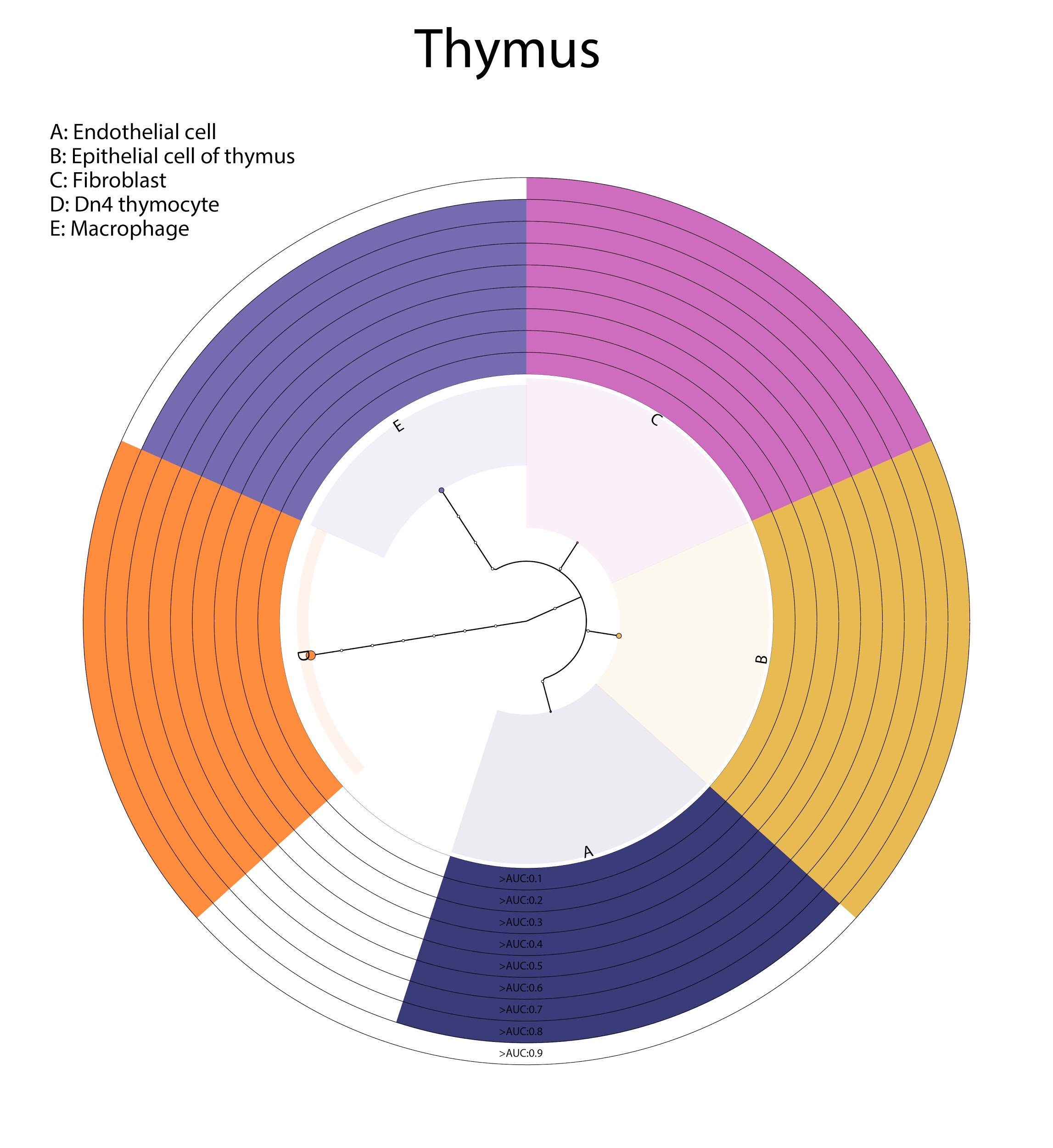

### **Supplementary Figure 21**

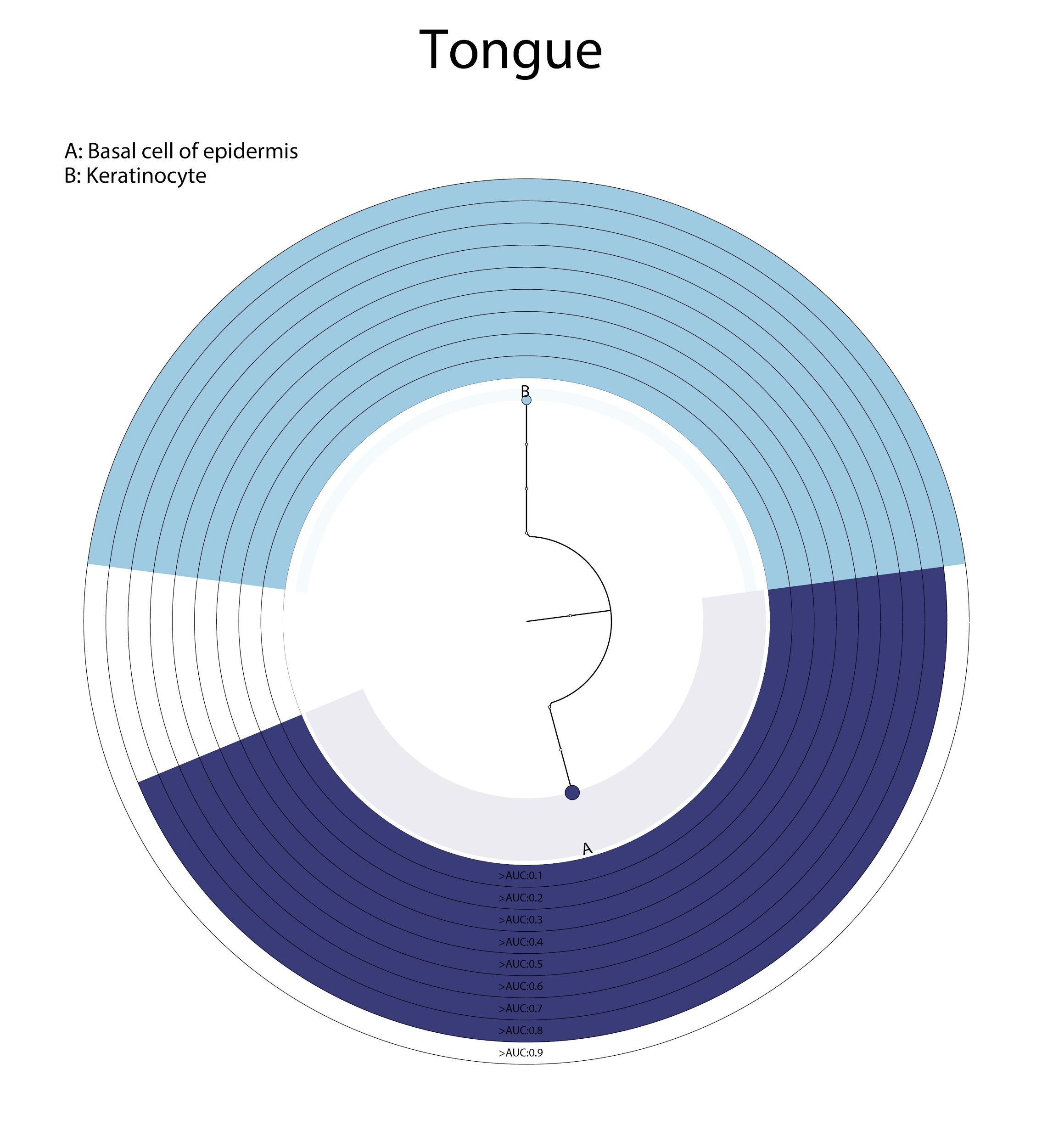

### **Supplementary Figure 22**

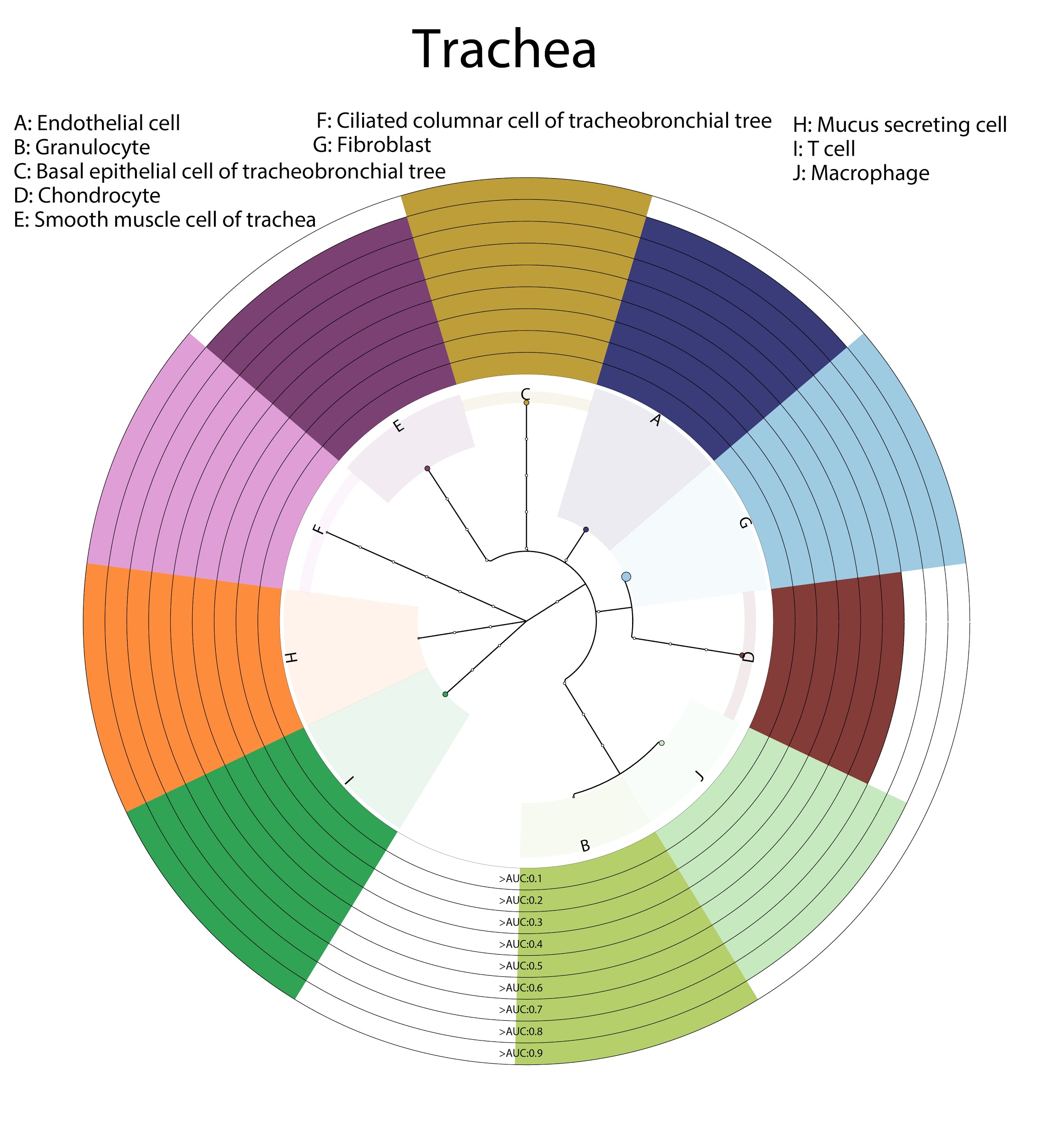

### **Supplementary Figure 23**
